# HuD is a neural enhancer of global translation acting on mTORC1-responsive genes and sponged by the Y3 small non-coding RNA

**DOI:** 10.1101/205658

**Authors:** Paola Zuccotti, Toma Tebaldi, Daniele Peroni, Marcel Köhn, Lisa Gasperini, Valentina Potrich, Tatiana Dudnakova, Guido Sanguinetti, Luciano Conti, Paolo Macchi, David Tollervey, Stefan Hüttelmaier, Alessandro Quattrone

## Abstract

The RNA-binding protein HuD promotes neurogenesis and favors recovery from peripheral axon injury. HuD interacts with many mRNAs, altering both stability and translation efficiency. UV-crosslinking and analysis of cDNA (CRAC) generated a nucleotide resolution map of the HuD RNA interactome in motor neuron-like cells. HuD target sites were identified in 1304 mRNAs, predominantly in the 3’UTR, with enrichment for genes involved in protein synthesis and axonogenesis. HuD bound many mRNAs encoding mTORC1-responsive ribosomal proteins and translation factors. Altered HuD expression correlated with the translational efficiency of these mRNAs and overall protein synthesis, in a mTORC1-independent fashion. The predominant HuD target was the abundant, small non-coding RNA Y3, which represented 70% of HuD interaction signal. Y3 functions as a molecular sponge for HuD, dynamically limiting its activity. These findings uncover an alternative route to the mTORC1 pathway for translational control in motor neurons that is tunable by a small non-coding RNA.

**Graphical abstract:** 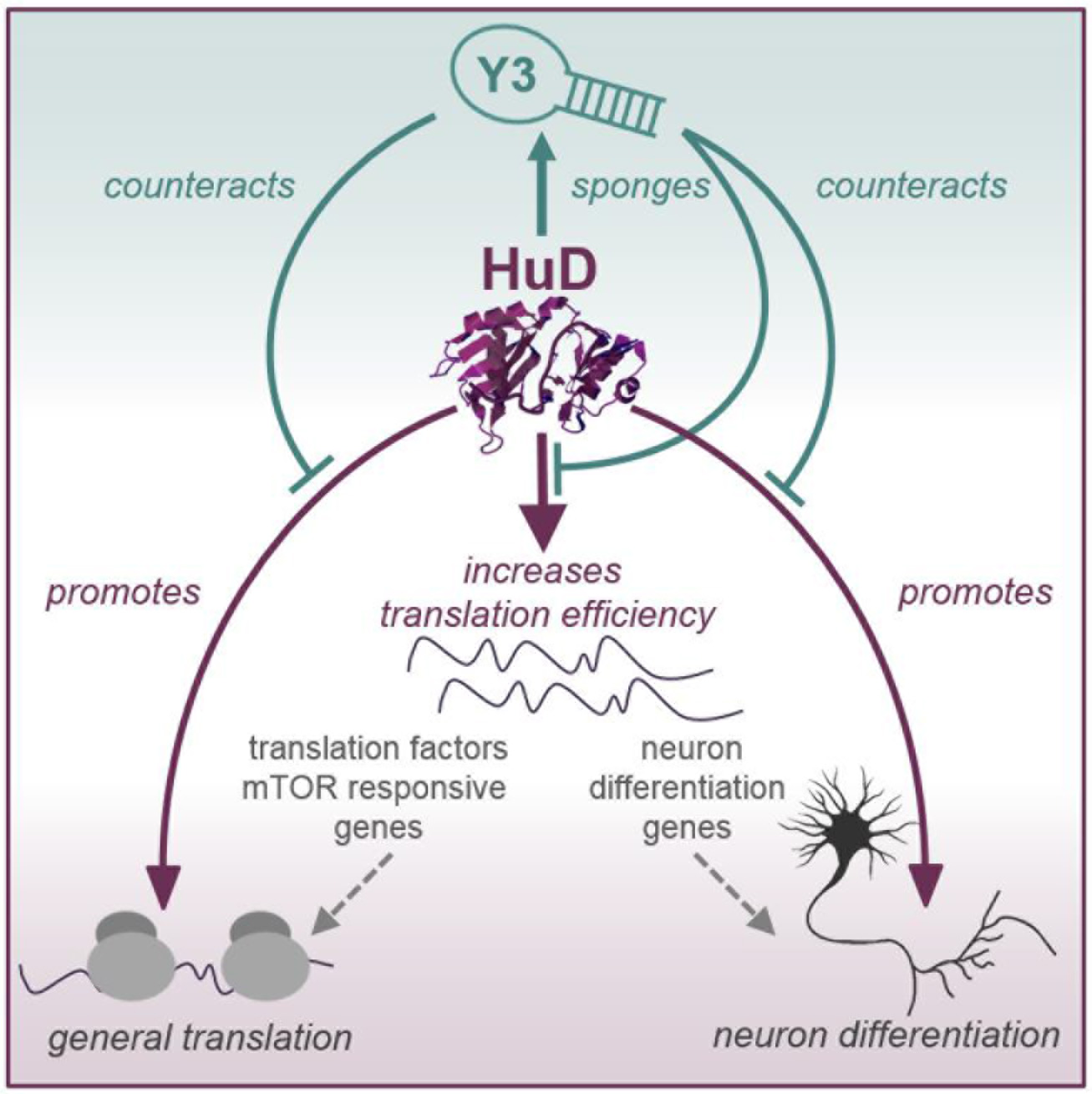

## Introduction

The intensively studied RNA-binding protein (RBP) <u>hu</u>man antigen <u>D</u> (HuD)/<u>e</u>mbryonic <u>l</u>ethal, abnormal <u>v</u>ision <u>l</u>ike 4 (ELAVL4) is predominately expressed in differentiated neurons, as are the other neuronal members (nELAV) of the ELAV family, HuB (ELAVL2) and HuC (ELAVL3). In contrast, HuR (ELAVL1) is ubiquitously expressed (Hinman and Lou, 2008; Pascale et al., 2008). HuD carries three <u>R</u>NA <u>r</u>ecognition <u>m</u>otif (RRM) domains and plays important roles in controlling the fate of many neuronal mRNAs (Bronicki and Jasmin, 2013). Functional analyses implicate HuD in the regulation of mRNA stability, alternative splicing, alternative polyadenylation, RNA localization and translation (reviewed in (Bronicki and Jasmin, 2013; Perrone-Bizzozero and Bird, 2013).

HuD is one of the first markers expressed during neuronal differentiation and plays a fundamental role in controlling neuronal cell fate. Loss of HuD induces increased self-renewal of the neural stem and progenitor cells (Akamatsu et al., 2005), whereas overexpression promotes neurite outgrowth, neurogenesis and neuronal plasticity (Marusich et al., 1994; Perrone-Bizzozero and Bolognani, 2002; Wakamatsu and Weston, 1997; Wang et al., 2015). HuD is activated by learning (Pascale et al., 2004; Quattrone et al., 2001) and HuD overexpression in transgenic mice leads to defects in hippocampal physiology and memory (Bolognani et al., 2007; Tanner et al., 2008).

Importantly, HuD is specifically implicated in motor neuron function and HuD knock-out mice show motor deficits, revealed by an abnormal hind-limb clasping reflex and a poor rotarod performance (Akamatsu et al., 2005). HuD levels in the superior cervical ganglion are highly reduced after axotomy, causing decrease of HuD target genes implicated in axonal regeneration (Deschenes-Furry et al., 2007), while regeneration following peripheral axon injury is associated with increased levels of HuD and of its direct target GAP43 (Anderson et al., 2003). In accordance with these functions, recent studies pointed out the intimate relationship between HuD and motor neuron diseases. HuD has been characterized for its ability to localize mRNAs in primary motor neurons and restore axon outgrowth defects in SMA motor neurons (Akten et al., 2011; Fallini et al., 2011, 2016). Moreover, cytoplasmic inclusions of TDP43, an almost universal pathological hallmark of Amyotrophic Lateral Sclerosis (ALS), are proposed to specifically sequester HuD (Fallini et al., 2012).

To understand the molecular mechanism that underpins the functions of HuD and the effects of its misregulation in neuronal physiology and pathology, we sought to positionally identify its RNA targets in a comprehensive way. Specific antibodies for individual nELAV paralogs are currently not available, so cross-linking and immunoprecipitation (CLIP) analysis (Konig et al., 2011; Scheckel et al., 2016) identified only RNAs cumulatively bound to nELAV proteins (HuB, HuC, and HuD). Specific HuD targets were previously identified by immunoprecipitating myc-tagged HuD from a HuD-overexpressing mouse strain (Bolognani et al., 2010). However, this approach does not provide information on the binding sites on RNA or distinguish between direct and indirect targets (Keene et al., 2006). To overcome these limitations, we specifically characterized the RNA interactome of HuD using the CRAC (cross-linking and analysis of cDNAs) method (Bohnsack et al., 2012; Granneman et al., 2009). This approach combines affinity purification of tagged HuD with the deep sequencing of directly bound RNA fragments. We performed our analysis in NSC-34 cells, which recapitulate motor neuron phenotypes *in vitro* (Cashman et al., 1992; Madji Hounoum et al., 2016).

We found that HuD directly and specifically enhances the translational efficiency of dozens of mRNAs known to be involved in motor neuron differentiation and axonogenesis, providing the molecular basis for its role in motor neuron development and physiology. Surprisingly, we also found that a major HuD-bound cluster contains mRNAs encoding components of the translational machinery. Our data show that HuD binding increases the translational efficiency of the bound mRNAs. This activity likely underlies the sustained promotion of translation we observed in motor neuron cells overexpressing HuD. This activity is also independent from the major pathway affecting general translation, controlled by the mTORC1 complex, despite targeting an overlapping set of mRNAs.

Remarkably, the Y3 small noncoding ncRNA (ncRNA) was by far the strongest HuD binding partner. Y RNAs are ncRNAs transcribed by RNA polymerase III (Köhn et al., 2013; Kowalski and Krude, 2015), ranging in size from 70 to 115 nucleotides and folding into characteristic stem-loop structures. There are four distinct Y RNAs in humans (hY1, hY3, hY4 and hY5) and only two Y RNAs in mice (Y1 and Y3). Y RNAs were proposed to be involved in efficient DNA replication and histone mRNA processing (Christov et al., 2006; Köhn et al., 2015). However, their biological functions are still largely elusive. Here, we demonstrate that Y3 acts as a molecular sponge for HuD activity.

## Results

### CRAC reveals the spectrum of HuD-bound RNAs in a motor neuron cell line

HuD shares a high sequence and structure similarity with the other members of the ELAV family, especially the nELAV HuB and HuC proteins, and all available antibodies fail to distinguish among them. This has been a major problem in the identification of specific HuD interactions. To overcome this difficulty and selectively identify *in vivo* HuD binding sites we adapted the CRAC protocol, originally developed for yeast, to be used with mouse motor neuron NSC-34 cells engineered with doxycycline-inducible His-HA tagged HuD. The experiment was performed in biological triplicate, with non-tagged but doxycycline-treated NSC-34 cells expressing only the tetracycline receptor (Trex cells) as control for non-specific signal (Figure 1A and **Experimental Procedures**). To precisely map the HuD RNA interactome, we developed and implemented a dedicated computational methodology (see also **Experimental Procedures**). This approach takes advantage of cross-linking induced mutations - primarily micro-deletions - in order to identify candidate binding sites with nucleotide resolution (Figure 1B). To increase specificity, we penalized locations with aligned reads and deletions in control experiments. For each of the remaining locations, we calculated a combined p-value based on a) the number of deletions, b) the number of aligned reads (coverage). We selected a set of 753 sequences surrounding locations with a combined p-value < 0.05 to build a Positional Weight Matrix (PWM). This was then used as a “seed” matrix to score all the other candidate binding sites (Figure 1C). To create the seed PWM, we defined a region spanning seven nucleotides around the deletion site. This size choice is based on previous crystallographic studies resolving the structure of the RRM1 and RRM2 domains of HuD bound to canonical AU rich elements (Wang et al., 2011), and on the fact that the larger region surrounding the deletion site did not show any sequence peculiarity, except for a higher frequency of A/U nucleotides over C/G (60% and 40% respectively, Figure S1A). The core of CRAC-defined, HuD binding site contains a triplet of U nucleotides, preceded by a non-U nucleotide with C>G>A in preferential order (Figure 1C).

**Figure 1.**
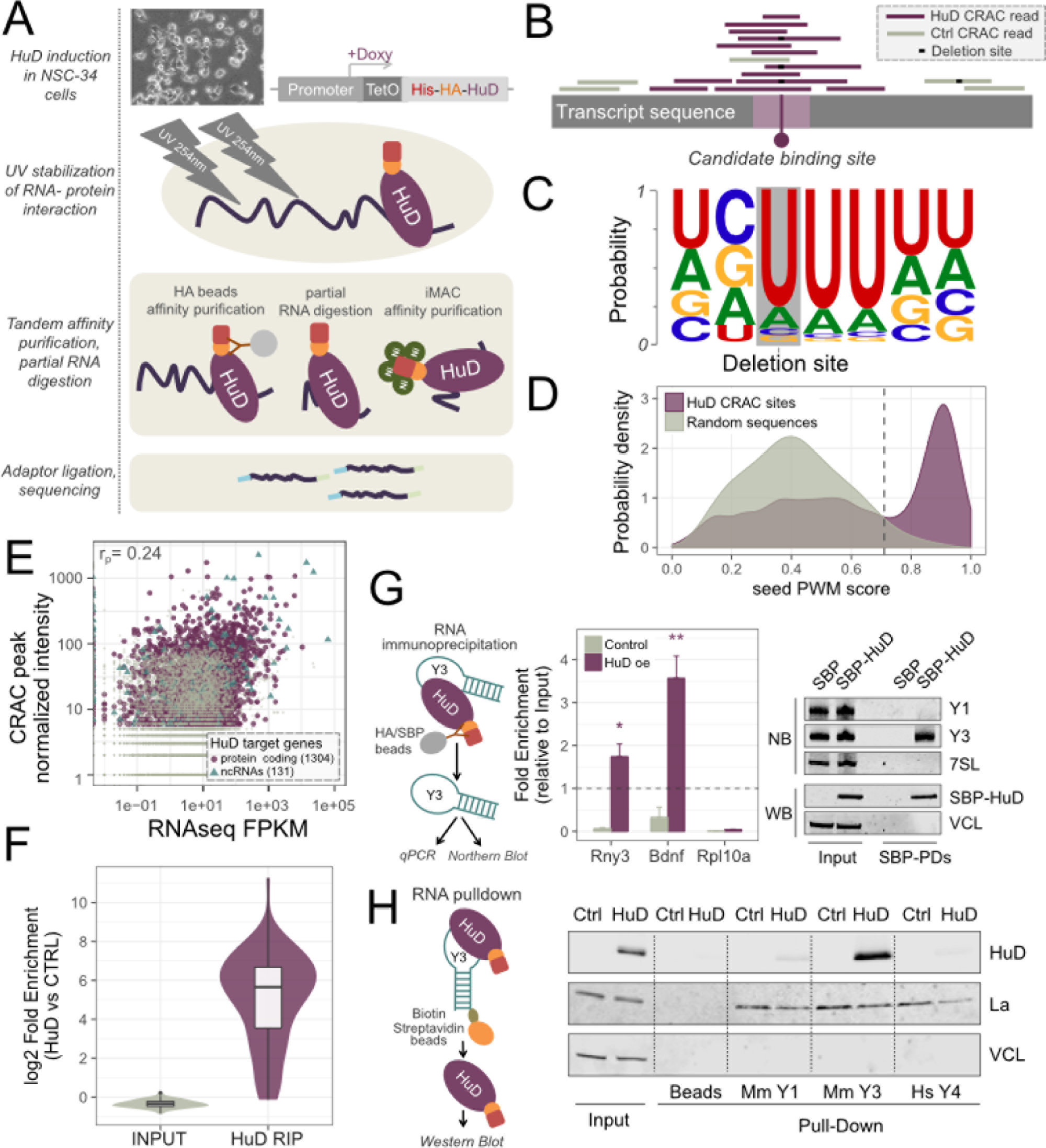
Defining the RNA interaction landscape of HuD in motor neuron cells. **(A)** Schematic representation of CRAC performed on motor neuron NSC-34 cells. **(B)** Overview of the computational methodology used to extract binding sites from CRAC raw data. **(C)** Logo representation of HuD seed PWM (Position Weighted Matrix) determined from unique CRAC cross-linking locations with combined Fisher p-value < 0.05. **(D)** Distribution of HuD PWM scores, calculated from CRAC deletion sites (in violet) and compared with random sequences (in grey). The score threshold to identify bona-fide binding sites was set as the 95th percentile of the random distribution (vertical dashed line). **(E)** Scatterplot displaying RNA-Seq transcript expression levels (FPKM) and normalized CRAC peak intensity values. The highest peak for each transcript is considered. HuD RNA targets are highlighted and annotated with their gene type. **(F)** Validation by RNA immunoprecipitation (RIP) and targeted sequencing of 70 HuD targets identified by CRAC. **(G)** Validation of HuD-Y3 interaction by alternative approaches: left panel, RNA immunoprecipitation followed by northern blots in HuD transfected NSC-34 cells. Transfection with mock vector was used as control; right panel, RNA immunoprecipitation followed by RT-qPCR in NSC-34 HuD inducible cells and in Trex NSC-34 cells (control); **(H)** Streptavidin pulldown of synthetic biotinylated Y RNAs (Y3, Y1 and human Y4) followed by Western blot analysis in NSC-34 cells induced for HuD expression. Trex NSC-34 cells were used as control. (In panels G, data are represented as mean ± SEM; t-test: *p < 0.05, **p < 0.01).

We used the seed PWM to score all deletion sites and select high-confidence HuD bound sites, with the advantage of selecting interaction sites even in transcripts with lower expression levels, for which the coverage was not enough to be included in the initial set, merely based on p-value. The strength of this methodology is shown by comparison of the distribution of scores associated with CRAC deletion sites with the distribution of random sequences (Figure 1D). The experimental distribution is peaked above the threshold score corresponding to the 95th percentile of the random distribution.

This approach detected 5,153 high confidence binding regions, mapped on 1,304 protein coding genes and 131 ncRNAs (Figure 1E, **Supplemental File 1**). Among the ncRNAs, there were 10 characterized long intergenic ncRNAs (lincRNAs), including Neat1, Malat1 and Yam1 that are known to be involved in cell-fate programming (Batista and Chang, 2013; Flynn and Chang, 2014). Strikingly, the predominant peaks of HuD binding were identified over the small ncRNA Y3 (Rny3), which represented 70% of all binding signal. Parallel RNA-Seq was performed in NSC-34 cells to quantify steady state transcript expression levels. For each transcript, the HuD binding site with the highest intensity was compared to RNA abundance. This comparison showed a low positive correlation between binding site intensities and transcript expression levels (measured by FPKM, pearson correlation = 0.24) (Figure 1E). Binding affinity should be therefore the main factor influencing peak intensity.

Interactions identified by CRAC were further validated by the RNA immunoprecipitation (RIP) technique for 70 protein coding genes and for the Y3 RNA. For the mRNAs tested, RIP followed by targeted sequencing confirmed the identification of *bona fide* HuD binding sites by CRAC, with a median log2 fold enrichment of 5.8 (Figure 1F). Interactions between HuD and Y3 were confirmed in the same cellular system. Y3 was selectively enriched together with the *Bdnf* mRNA (positive control) in HuD ribonucleoprotein particles, but not in the control cells (Figure 1G, left panel). For both conditions, no binding to the *Rpl10a* transcript (negative control) was detected. To avoid the possible involvement of the HA-His tags in the binding, we performed a second RIP in NSC-34 transiently transfected with SBP-tagged HuD. We checked the presence of Y3 and Y1, the only other member of the Y RNA family in the mouse genome, among the HuD bound RNAs, and confirmed also under this experimental condition the interaction between HuD and Y3. No binding was detected for Y1 RNA, which is structurally related to Y3, or the highly abundant small ncRNA signal recognition particle RNA (7SL) (Figure 1G, left lower panel). As a further validation, we performed a pulldown assay by using synthetic biotinylated Y RNAs Y3, Y1 and human Y4 (negative control), in both NSC-34 cells induced for HuD expression and control cells. Western blotting demonstrated specific association between HuD and Y3 (Figure 1H, right panel). La protein and Vinculin (VCL) were used as positive and negative controls, respectively, for Y RNA interacting proteins (Stefano, 1984).

### HuD enhances translation of translation factors

To provide a functional characterization of HuD-interacting RNAs, we performed enrichment analysis of Gene Ontology (GO) terms and pathways (KEGG and Reactome) (Figure 2A). Enrichment was seen for terms related to genes involved in mRNA processing and function; e.g. mRNA binding, splicing and localization. In addition, we found a strong enrichment (2.7 fold enrichment and 2.7E-10 FDR) for genes involved in translation: 80 genes, including 34 ribosomal components and 12 translation initiation or elongation factors. Within mRNA targets, HuD binding sites were predominantly located in the 3'UTR of protein coding transcripts (92% of mRNA binding sites), consistent with functions in translation (Figure 2B). However, there was no clear overall preference for regions proximal to either the translation termination site or the polyadenylation and cleavage site within the 3’UTR.

**Figure 2.**
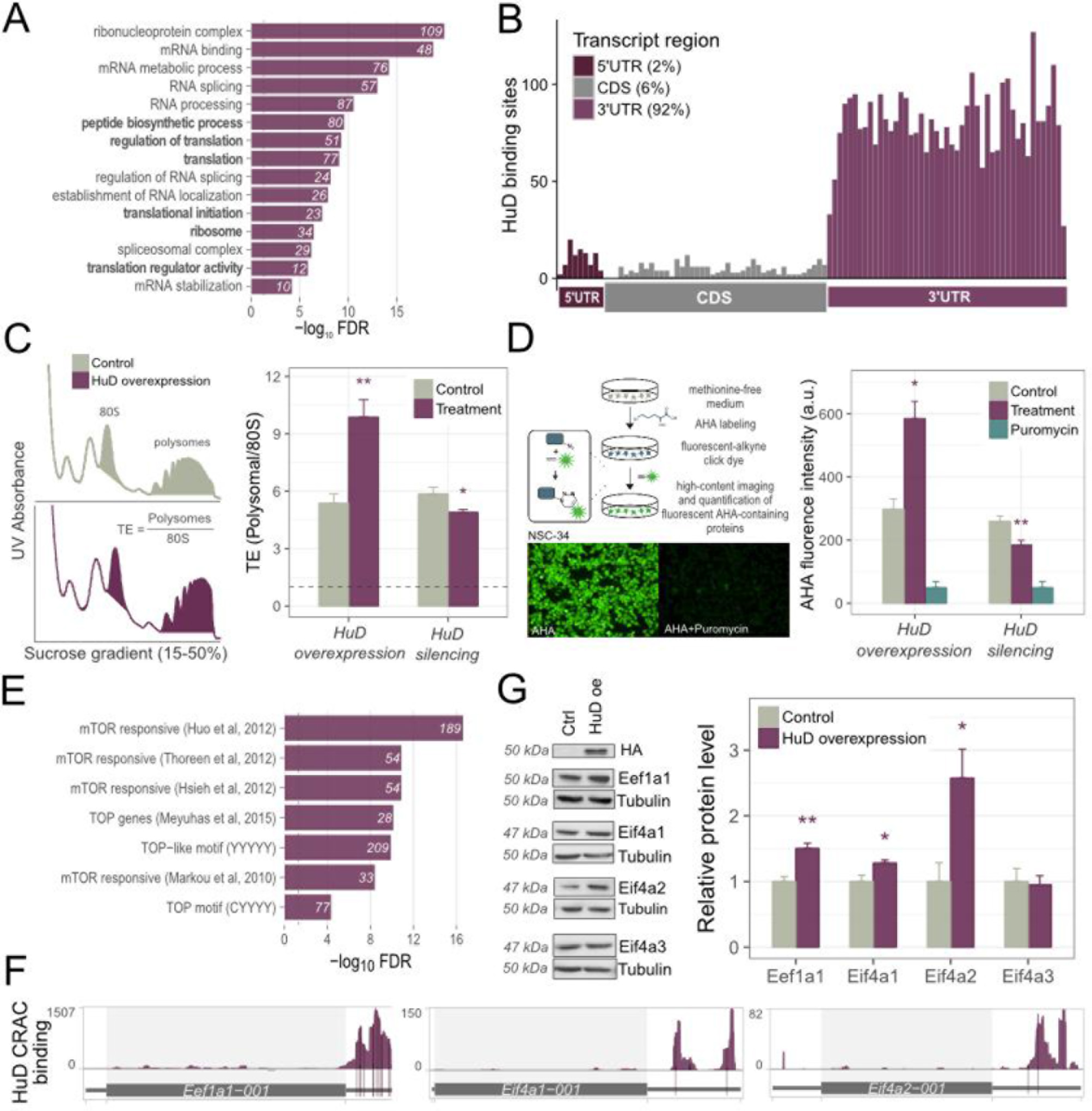
HuD increases global and target-specific translation. **(A)** Top enriched Gene Ontology terms among HuD mRNA targets are related to RNA processes, including splicing, transport, stability and translation (highlighted in bold). **(B)** Metaprofile of HuD binding sites along protein coding transcripts, showing binding enrichment in 3’UTRs. **(C)** Right panel: representative sucrose gradient profiles in control and HuD overexpressing NSC-34 cells. The translation output is calculated as the ratio between the area under polysome peaks and the area under the 80S peak. Left panel: calculation of the global translational output from sucrose gradient profiles upon HuD silencing and overexpression. **(D)** Right: schematic representation of Click-IT^®^ AHA (L-Azidohomoalanine) chemistry assay to quantify de-novo protein synthesis in NSC-34 cells. Left: detection of de novo protein synthesis by AHA upon HuD silencing and overexpression. Puromycin, a translation inhibitor, was used as negative control. Acquisition and automated quantification was performed by high content screening device. **(E)** Enrichment of mTOR responsive mRNAs among HuD targets, as listed in multiple literature sources. **(F)** Positional binding maps for three mTOR responsive HuD targets: Eef1a1, Eif4a1 and Eif4a2. **(G)** Western blot analysis of HuD targets (Eef1a1, Eif4a1, Eif4a2) and negative control (Eif4a3) in HEK-293 cells transiently transfected with HuD, relative to transient transfection with an empty vector. Tubulin was used as reference gene. (In panels C,D,G, data are represented as mean ± SEM; t-test: *p < 0.05, **p < 0.01).

The preferential binding to mRNAs encoding ribosomal proteins and translation factors suggested that HuD could promote global translation through the post-transcriptional modulation of these mRNAs. We therefore assessed the role of HuD in modulating global translation by polysome profiling in NSC-34 cells with overexpression of the inducible His-HA HuD. The global translational efficiency (TE) of the cells was calculated as the ratio between the absorbance of polysomes and the total absorbance of non-translating 80S ribosomes (see **Experimental Procedures** and Figure 2C). As shown in Figure 2C, HuD overexpression significantly increased the global TE of NSC-34 cells. Conversely, HuD depletion by RNA interference resulted in a reduced global TE. To support this finding we assessed the ability of HuD to promote global nascent translation. *De novo* protein synthesis was quantified using metabolic labelling, based on the incorporation of the methionine analogue L-azidohomoalanine (AHA) into newly synthesized proteins. AHA was added to HuD overexpressing cells or to HuD silenced cells and we used click-chemistry to fluorescently tag and quantify the incorporated AHA. A significant increase in overall *de novo* protein synthesis was detected in HuD-overexpressing NSC-34 cells compared to control, whereas the knockdown of HuD resulted in *de novo* protein synthesis reduction (Figure 2D). As a control for non-specific signals, the analysis was also performed in the presence of the translation inhibitor puromycin.

To investigate the mechanism through which HuD promotes global translation, we specifically focused on translation-related HuD mRNA targets identified by the CRAC analysis. Notably, many of these factors are known to be mTOR responsive (Hsieh et al., 2012; Larsson et al., 2012; Thoreen et al., 2012), including 5'-TOP or 5'-TOP-like mRNAs (Meyuhas and Kahan, 2015) (Figure 2E). Among these mRNAs, we selected for further investigation the translation elongation factor *Eef1a1*, which contains a classical terminal oligopyrimidine (TOP) motif in the 5’UTR (Levy et al., 1991; Meyuhas and Kahan, 2015) and the eukaryotic initiation factors *Eif4a1* and *Eif4a2*, coding for the DEAD-box helicases belonging to the eiF4F complex that act as strong stimulators of cap-dependent translation by 5’UTR mRNA unwinding. *Eef1a1*, *Eif4a1* and *Eif4a2* mRNAs are strongly bound by HuD in their 3’UTRs (Figure 2F). As shown in Figure 2G, overexpression of HuD significantly increased the protein levels of Eef1a1, Eif4a1 and Eif4a2 with respect to tubulin. As negative control we used the exon junction complex component Eif4a3 (Chan et al., 2004), which is a Eif4a1 and Eif4a2 paralogue but is not an identified HuD target or known to be involved in translation. Levels of Eif4a3 were unaffected by enhanced HuD expression.

Since mTOR responsive genes were significantly enriched among HuD targets (Figure 2D), we next assessed if the HuD-dependent boost to global and target specific translation was mediated through the mTORC1 pathway. To address this question, cells were serum starved for 8h to decrease activity of the mTORC1 pathway to less than 50%, as assessed by the phosphorylation status of its two terminal target proteins Eif4e-bp1 and Rps6. This treatment did not affect the levels of endogenous HuD or inducible His-HA-HuD (Figure 3A), and did not induce P-bodies or stress granules (Figure S2A) in NSC-34 cells. Global TE was then measured by polysome profiling. As expected, starvation caused a decrease in the global TE compared to serum repleted cells. Interestingly, HuD overexpression restored and even increased TE in serum depleted cells relative to repleted cells (Figure 3B). The effects of the mTORC1 inhibitor Torin1 were also suppressed by HuD overexpression (Figure S2B). Variations in the TE of specific mRNAs were determined from the ratio between changes in polysomal and total RNA. We selected different classes of mTOR-responsive, HuD bound mRNAs for TE quantification by qPCR: ribosomal proteins, Pabpc1, elongation factors and initiation factors. The results consistently showed that HuD overexpression increased the TE of the target mRNAs upon starvation (Figure 3C), whereas Eif4a3, which is not bound by HuD, was unaffected. The TE changes of HuD targets consistently resulted from increased representation in the polysomal RNA fraction rather than reduction in total RNA. We further verified that the TE increase correlated with an enhanced Eef1a1 protein level, with no effect on the negative control Eif4a3 (Figure 3D).

**Figure 3.**
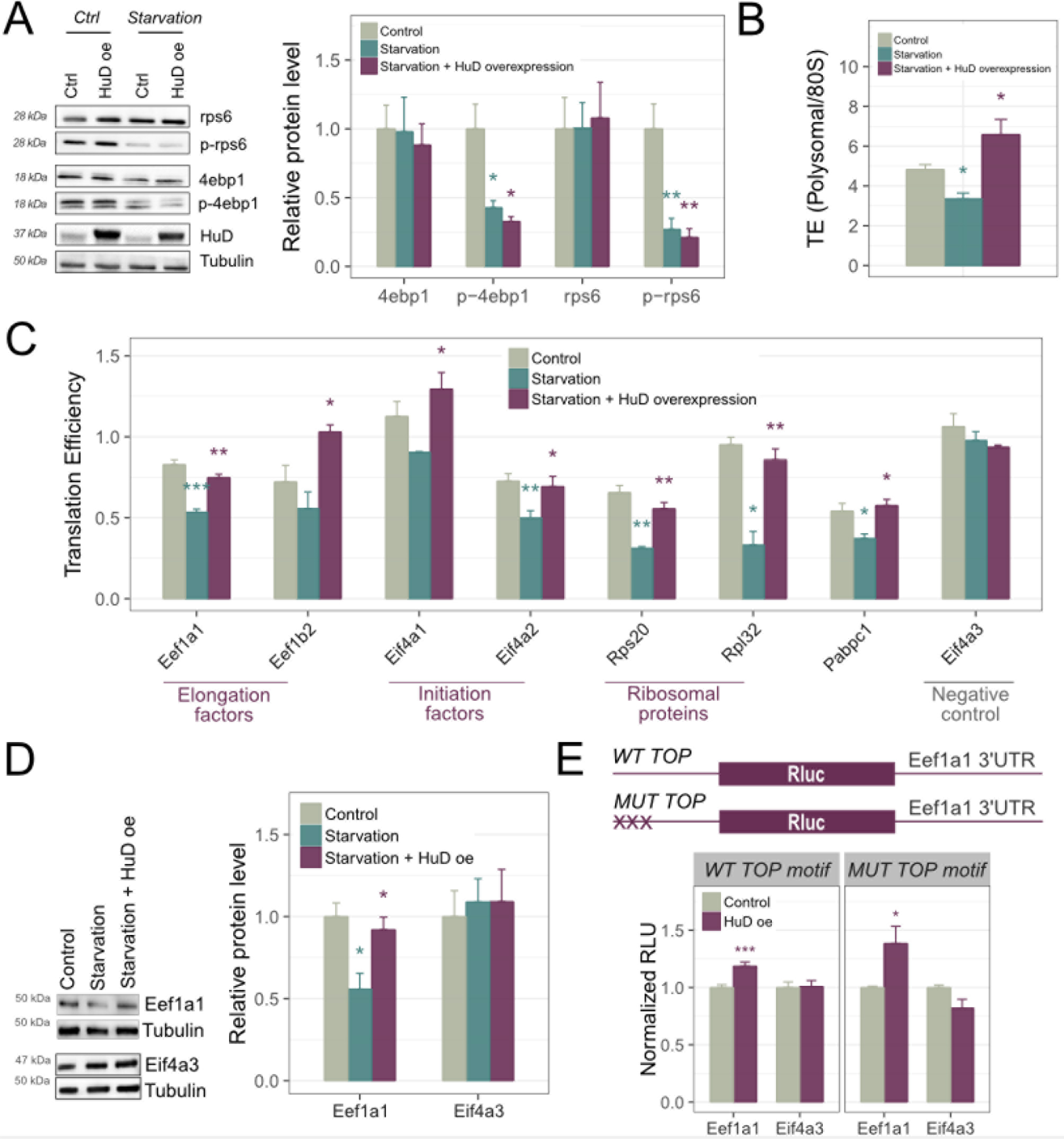
HuD enhancement of global and target specific translation efficiency does not depend on the mTORC1 pathway. **(A)** Left, western blot analysis of Rps6 and Eif4ebp1 phosphorylation following serum-deprivation (8 hours) in NSC-34 cells **(B)** Measurement of global TE by sucrose gradient centrifugation in the following conditions: control, starvation and starvation coupled with HuD overexpression. Global TE is expressed as the ratio between the area under polysome peaks and the area under the 80S peak. **(C)** TE quantification of selected mTOR-responsive mRNAs in control, starvation and starvation coupled with HuD overexpression conditions. Target-specific TE represents the ratio between polysomal and total RNA changes measured by RT-qPCR in NSC-34 cells. Gapdh and Als2 were used as reference genes. **(D)** Western blot analysis of Eef1a1 and Eif4a3 in NSC-34 cells collected in three different conditions: control, starvation and starvation with HuD overexpression. **(E)** Barplot displaying normalized luciferase intensity values in HEK-293 cells transiently transfected with HuD, relative to transient transfection of the empty vector. Cells were co-transfected with wild type (WT) or mutated (MUT) TOP motif bearing luciferase vectors with the 3’UTR of Eef1a1 (HuD target) or Eif4a3 (negative control). (In panels A,B,C,D,E, data are represented as mean ± SEM.t-test *p < 0.05, **p < 0.01 and ***p < 0.001. In panels A,B,C, “Starvation” was compared to “Control”, and “Starvation + HuD overexpression” was compared to “Starvation” for testing statistical significance).

These results indicated that HuD is able to rescue global and target-specific translation inhibition when the mTORC1 pathway is inhibited. To confirm this, we further explored how HuD regulates the expression of the 5’ TOP gene Eef1a1, known to be selectively modulated by mTORC1 (Thoreen et al., 2012). We cloned the 3’ UTRs of *Eef1a1* and the negative control *Eif4a3* downstream from luciferase, in a reporter vector harboring an ectopic canonical TOP motif at the 5’ end (Thoreen et al., 2012). We expressed these reporters alone or in combination with HuD in HEK-293 cells. Luciferase activity was enhanced by HuD co-expression for the vector carrying the 3’UTR from *Eef1a1*, but not for *Eif4a3* (Figure 3E). We conclude that HuD enhances translation of mRNAs bearing TOP motifs in the 5’UTR only if there is an HuD binding site in the 3’UTR. To determine whether HuD requires a TOP motif to enhance translation, we used a luciferase vector bearing a mutated TOP motif not responding to mTOR signaling (MUT-TOP) in the 5’UTR (Figure 3E). HuD overexpression increased the luciferase activity elicited by wild type or mutated TOP-bearing luciferase vectors with the *Eef1a1* 3’UTR, while no increase was detected in constructs containing the *Eif4a3* 3’UTR (Figure 3E). These results collectively demonstrate that the translational control exerted by the mTORC1 pathway on 5’ UTR TOP mRNAs can be tuned by the independent translational enhancement promoted by HuD through 3’UTR binding.

### HuD stimulates translation of mRNAs involved in neuronal fate commitment and in axonogenesis

The control of translation is a key step in mediating neuronal plasticity. Particularly in motor neurons, fine tuning of protein synthesis in distinct cell compartments (soma, dendrites and axons) is required to modulate neuronal activity and synaptic plasticity (Holt and Schuman, 2013; Jung et al., 2012). HuD was demonstrated to act on specific target mRNAs involved in neuronal differentiation in cell line models (Deschenes-Furry et al., 2007). We identified as high confidence hits multiple neuronal mRNAs previously reported to interact with HuD in the HuD CRAC data (**Supplemental File 1**).These included App (Kang et al., 2014), Atg5 (Kim et al., 2014), Gls (Ince-Dunn et al., 2012), Ikzf5 (Bellavia et al., 2007), Lmo4 (Chen et al., 2007), Marcks (Wein et al., 2003), Msi1 (Ratti et al., 2006), Nova1 (Ratti et al., 2008), Nrn1 (Akten et al., 2011; Tiruchinapalli et al., 2008; Wang et al., 2011).

Analysis of mRNAs responsible for neuronal specification and identity in the CRAC data revealed enrichment for two partially overlapping classes (20% superimposition): genes involved in neuronal differentiation and neurogenesis (135 genes), and genes more specifically annotated as linked to axonogenesis, axon guidance, myelin deposition, axon localization and synaptic functionality (112 genes) (Figure 4A). To assess whether HuD binding to these mRNAs results in phenotypic effects on neurogenesis, we induced HuD overexpression in differentiating NSC-34 cells. Multiple morphological parameters of differentiation were measured, including the number of neuronal extremities, roots and segments. We observed a significant increase in neuronal outgrowth and branching in HuD overexpressing cells compared to control cells (Figure 4B **and** Figure S3A). Therefore, we inspected whether HuD expression correlated with enhanced TE for 11 selected HuD target mRNAs, known to play important roles in motor neurons and axons (Hua et al., 2015; Martinez et al., 2016; Song et al., 2009; Wu et al., 2017). As shown in Figure 4C, we found a significant TE increase in HuD overexpressing cells for each of these mRNAs. The increases were greater for Kif5b, Sema4d, Picalm, Acsl4 and Hnrnpa2b1. As above, TE enhancement upon HuD overexpression was driven by increased polysomal occupancy, with almost no variation in total RNA levels.

**Figure 4.**
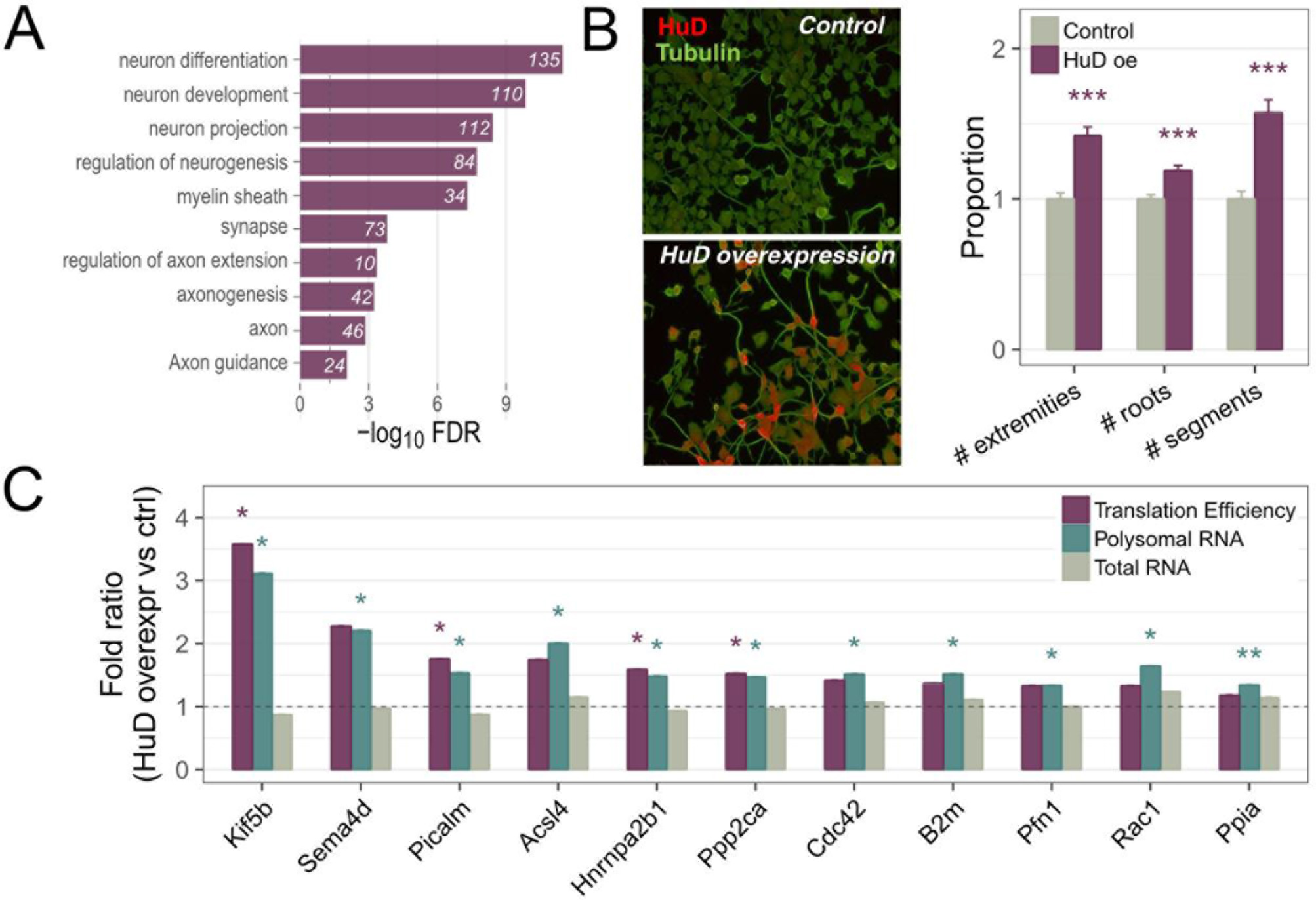
HuD increases the TE of several neuronal-related targets, supporting neuronal differentiation. **(A)** Gene ontology neuronal-related enriched terms among HuD targets. **(B)** Representative images of differentiated NSC-34 cells (control or overexpressing HA-tagged HuD) immunostained with anti-HA tag (red) and anti-tubulin antibodies (green) (left); Neurite outgrowth quantification in differentiated NSC-34 cells overexpressing HA-tagged HuD compared to control. Multiple parameters (number of neuronal extremities, roots and segments) were analyzed using High content screening (HCS) (right). **(C)** Measurement of TE variations on specific neuronal-related targets in HuD overexpressing cells compared to control. TE was calculated as the ratio between the polysomal and total RNA quantified by targeted sequencing and displayed in the plot in green and grey, respectively. (In panels B,C data are represented as mean ± SEM. t-test *p < 0.05, **p < 0.01 and ***p < 0.001).

HuD was shown to restore axon outgrowth and protein localization in SMA motor neurons (Fallini et al., 2016). We therefore examined the overlap between HuD binding targets and mRNAs with altered expression in the motor neuron disease state, previously determined in transcriptome-wide studies (Figure S3B). Strong enrichment was observed for motor neuron disease-associated genes among HuD targets. The effects of HuD overexpression on translation was determined for specific genes associated with ALS (Vcp, Ubqln2, Pfn1), and genes with altered expression in both ALS and spinal muscular atrophy (SMA) (Figure S3C). This observation highlights a potential role for HuD in modulating the expression of pathologically relevant transcripts in motor neurons.

### HuD is sequestered by the Y3 ncRNA

Quantitative analysis of HuD CRAC interactions by peak intensity clearly identified the 102 nt ncRNA Y3 as the largely dominant target. Inspection of the CRAC deletion profiles identified two binding sites in Y3 that map to loop regions, which are closely positioned in the secondary structure (Teunissen et al., 2000) (Figure 5A). Both these binding sites contain the “VUUU” consensus identified by CRAC as the sequence motif preferentially bound by HuD.

**Figure 5.**
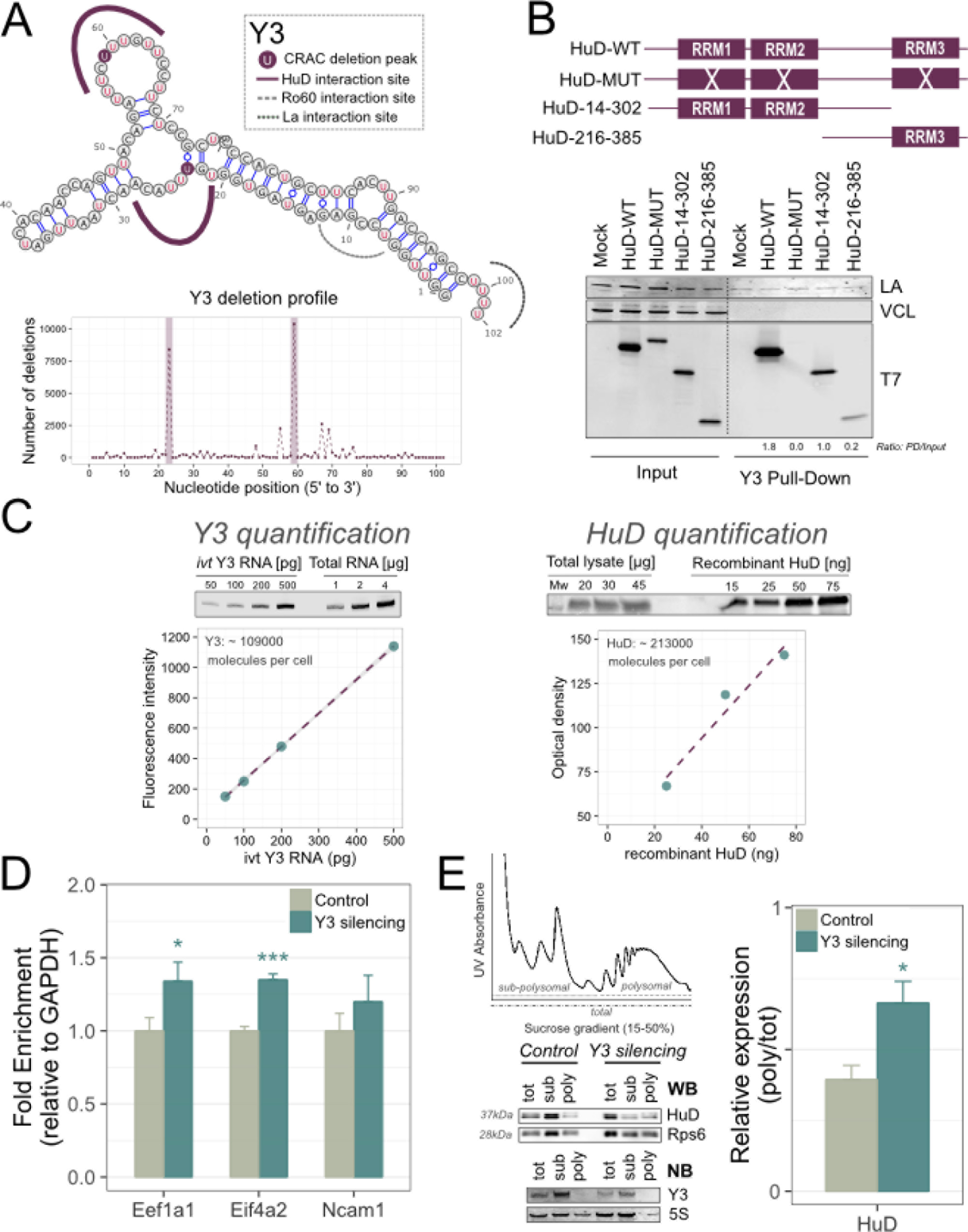
Y3 reduces HuD association with target RNAs and polysomes. **(A)** Upper panel: secondary structure of Y3 with HuD interaction sites (visualized with VARNA) based on chemical probing. Lower panel: based on CRAC deletion profiles, HuD binds to Y3 in two discrete locations, both positioned in polypirimidine tracts in the looped region outside the Y3 structural stem. **(B)** Y3 RNA-pulldown showing that HuD interacts with Y3 by the RRM domains, mainly RRM1 and RRM2. **(C)** Quantification of Y3 and HuD molecules number in NSC-34 cells. The estimated molecule number was calculated by mean of a calibration plot generated by known amounts of standards, i.e. In vitro transcribed (ivt) Y3 RNA and recombinant HuD, respectively. **(D)** RNA immunoprecipitation analysis of HuD binding to Eef1a1, Eif4a2 and Ncam1 mRNAs after Y3 silencing; data were normalized to control Gapdh mRNA levels in each IP. **(E)** Polysomal versus total localization of HuD in Y3 depleted NSC-34 cells, compared to control cells. HuD levels were measured by Western blot, while Y3 levels were measured by Northern Blot. (In panels D and E data are represented as mean ± SEM. t-test *p < 0.05, ***p < 0.001).

To determine the region of HuD involved in Y3 binding, we transiently transfected NSC-34 cells with four different HuD constructs (Fukao et al., 2009): i) wild type (wt), ii) HuD-MUT, lacking any RNA-binding activity, iii) HuD-14-302, lacking RRM3, the HuD RNA binding domain proposed to bind the polyA, and iv) HuD-216-385, lacking the RNA binding domains RRM1 and RRM2. Subsequently, we performed Y3 pull-down to define the HuD regions responsible for the binding. We found that the HuD RRM domains are necessary for the interaction with Y3, as no binding can be detected for HuD-MUT, with a stronger contribution of the first and the second RRMs (Figure 5B).

To investigate whether Y3 could influence HuD expression levels or vice versa, we silenced either Y3 and HuD expression by siRNA, monitoring the expression levels of HuD and Y3, respectively. As we did not report any changes, we concluded that the interaction between HuD and Y3 does not clearly influence their expression levels (Figure S4).

We hypothesized that the high levels of Y3 binding might compete with HuD interactions on mRNA, qualifying Y3 as a molecular decoy or “sponge” (Köhn et al., 2013) for HuD. Using calibration curves, we estimated that NSC-34 cells contain on average approximately 213000 molecules of HuD protein and 109000 molecules of Y3 RNA (Figure 5C). If the two HuD binding sites on Y3 are occupied by different HuD molecules, this estimated ratio (1.95) suggested that Y3 might be able to potentially sequester much or all of the HuD population. To test this hypothesis, we first evaluated whether Y3 could compete HuD binding to other RNA targets. HuD was immunoprecipitated from NSC-34 cells with or without prior treatment with siRNAs directed against Y3, and three HuD-associated mRNAs (*Eef1a1*, *Eif4a2* and *Ncam1*) were quantified. Cells depleted for Y3 showed increased co-precipitation with HuD for all three targets **(Figure 5D)**. This finding supports competition for HuD binding between mRNAs and Y3.

HuD can dynamically associate with polysomes (Atlas et al., 2007; Bolognani et al., 2004). To determine whether Y3 affects this activity, we isolated polysomes from NSC-34 treated or not with Y3 siRNAs. Evaluation of HuD recovery in the polysomal fraction relative to the total cellular protein showed increased polysome association after Y3 depletion (Figure 5E). We conclude that Y3 can sequester HuD away from mRNAs and the translation machinery.

To assess the functional consequence of HuD sequestration by Y3, we tested whether Y3 modulates translation enhancement by HuD. After Y3 depletion in NSC-34 cells (Figure 6A), we assessed both the global translational output and *de novo* translation. Measurement of the TE, by the ratio of polysomes and the 80S monosomes following gradient fractionation, indicated increased ribosome engagement in active translation (Figure 6A) and this was supported by increased AHA incorporation following Y3 depletion (Figure 6B). These results indicate that Y3 acts as a general repressor of translation in NSC-34 cells.

**Figure 6.**
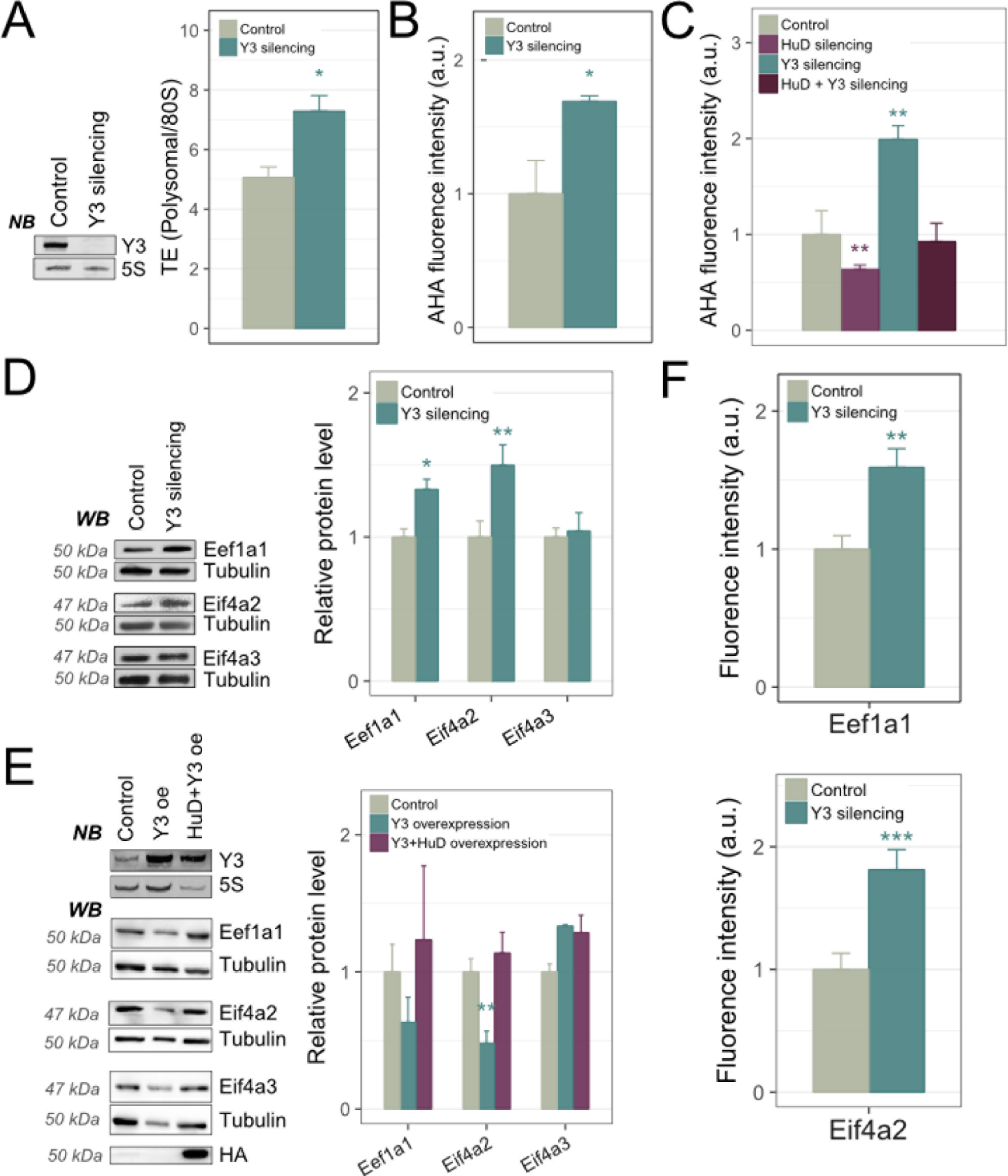
Y3 modulates HuD functions and targets. **(A)** Global translation output by sucrose gradient profiles upon Y3 silencing in NSC-34 cells. **(B)** De novo protein synthesis by AHA labeling upon Y3 silencing in NSC-34 cells. **(C)** AHA labeling experiments in NSC-34 cells depleted for Y3, for HuD or for both, showing antagonism between Y3 and HuD on protein synthesis. **(D)** Western blot of HuD targets (Eef1a1, Eif4a2) and negative controls (Eif4a3) in NSC-34 cells transiently silenced for Y3, relative to control silencing. **(E)** Quantification of Eef1a1 and Eif4a2 protein levels in primary motor neurons transfected with an shRNA construct directed against Y3 (sh_Y3) or a control vector (sh_Ctrl) (In panels A, B, C, D, E, F data are represented as mean ± SEM. t-test *p < 0.05, **p < 0.01 and ***p < 0.001).

To determine the relation between HuD and Y3 in the modulation of translation, we measured global de novo translation by AHA incorporation in NSC-34 cells after the following treatments: (i) HuD silencing by siRNA, (ii) Y3 depletion by shRNA expression, (iii) combined silencing of HuD and Y3. Co-depletion of Y3 partially restored the reduction in translation seen following knock down of HuD alone (Figure 6C). These data indicate that the impact of HuD silencing on translation is mitigated if HuD sequestration by Y3 depletion is also reduced, presumably due to an increase in the available pool of HuD **(Figure 6C)**. To prove that the Y3 modulatory effect on global translation is at least partially mediated by the altered expression of HuD targets, we depleted Y3 in NSC-34 cells and assessed the protein levels of Eef1a1, Eif4a2 and the negative control Eif4a3. We observed a significant increase for both HuD targets, but not for the negative control **(Figure 6D)**. Hence, Y3-mediated regulation of global translation is at least partly due to its modulation of HuD target genes. We also tested the proposed molecular competition between Y3 and HuD by ectopic expression in HEK-293 cells **(Figure 6E)**. Overexpression of Y3 was associated with specific decrease in the protein levels of EEF1A1 and EIF4A2, whereas no change was observed for the negative control EIF4A3. Co-expression of HuD restored protein expression of EEF1A1 and EIF4A2 to control levels.

Functional interactions between Y3 and HuD were further analyzed in primary cultured embryonic motor neurons (MNs). As previously reported (Fallini et al., 2011), HuD displays a distinctive granular pattern of localization in MNs (Figures S5A, S5B). Notably, MNs have high level of endogenous Y3, mainly localized to the axonal compartment (Figures S5C, S5D). To test the effect of Y3 depletion on HuD targets in MNs, we performed transfection with either an shRNA vector targeting Y3 (shY3) or the empty control. The silencing efficiency of the shY3 vector was tested (Figure S5E), and the co-expression of the green fluorescent protein (GFP) allowed us to identify silenced MNs. Three days after transfection, MNs were fixed and immunostained for Eef1a1 and Eif4a2. Compared to control cells, shY3-treated MNs showed a significant increase in Eef1a1 and Eif4a2 protein levels, recapitulating the data obtained in NSC-34 cells (Figure 6F **and S5F)**.

### Y3 blocks the function of HuD in neuronal differentiation

HuD has an established role in promotion neuronal differentiation. It seemed possible that developmentally regulated switch in HuD:Y3 ratio might control HuD availability for activity on mRNA targets, thus boosting neuronal differentiation in a specific temporal window. We initially analyzed changes in HuD and Y3 levels and ratio during neuronal development. To test this, we took converted mouse Embryonic Stem Cells (mESCs) into neurons using a defined neuronal differentiation protocol (Ying et al., 2003). Under these conditions, mESCs are efficiently committed to the neural progenitor lineage and subsequently begin a neurogenic phase that leads to the differentiation into neurons. We measured HuD and Y3 levels at three different stages of the differentiation procedure: mESC (D0), neural progenitors (D7) and early neurons (D10). There was progressive increase in levels of both Y3 and HuD during this process, but with different kinetics (Figure 7A). Y3 showed a substantial increase at the neural progenitor stage (2.5 fold at D7 relative to D0), but then showed only a modest further increase (3 fold at D10 relative to D0). In contrast, HuD exhibited a 5 fold increase at the neural progenitor stage (D7) and a 10 fold increase at the early neuron stage (D10).These results predict that a strong reduction in HuD sequestration by Y3 at the neurogenic stage *in vivo*, allows HuD to progressively drive neuronal differentiation.

**Figure 7.**
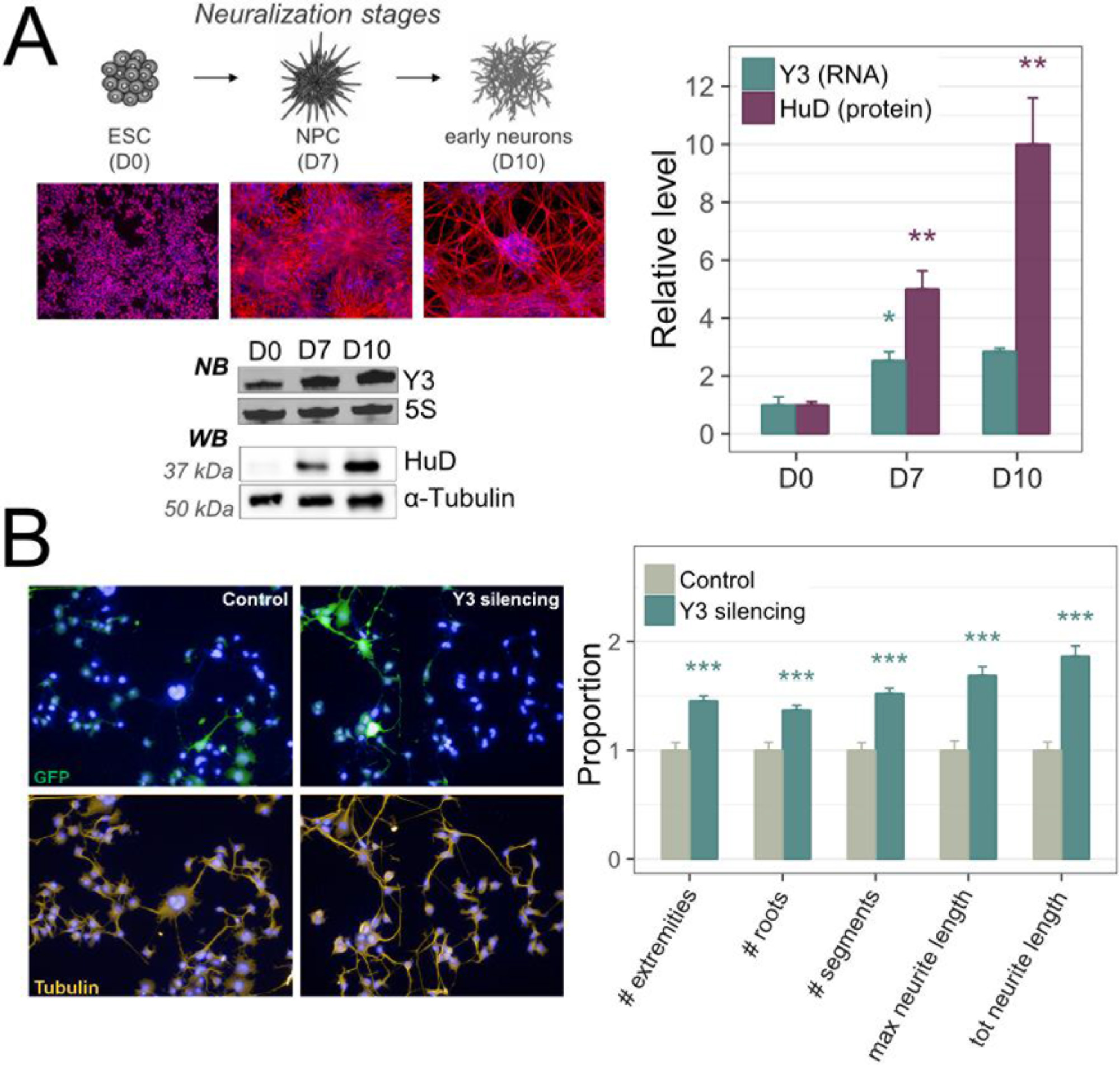
HuD and Y3 relative expression levels change along neuronal differentiation. **(A)** Differentiating ESCs cultures assayed for Y3 and HuD expression levels by northern blot and western blot, respectively. Cultures were fixed and immunostained for stage-specific markers: Oct4 (ESCs; red), Nestin (NPCs; red), beta3-tubulin (early neurons; red). Relative quantification of Y3 and HuD levels are shown (right). **(B)** Differentiated NSC-34 cells (control or silenced for Y3) immunostained with anti-tubulin antibody (yellow) to detect neurites (left panel); GFP (green) identified transfected cells that were subjected to high content analysis. Multiple parameters of neurite morphologies were analysed using Operetta HCS device (right panel). (In panels A, B data are represented as mean ± SEM. t-test *p < 0.05, **p < 0.01 and ***p < 0.001).

To directly test for a negative role for Y3 in neuronal differentiation, we induced shY3 expression in NSC-34 cells exposed to differentiation conditions. Y3 depletion significantly increased neurite extension and branching in comparison to control NSC-34 cells (Figure 7B).

Taken together, our results show that Y3 directly binds to HuD, sequestering it from target mRNAs and counteracting its ability to promote translation and neuronal differentiation.

## Discussion

The crucial role of HuD in motor neuron plasticity and axon regeneration (Akamatsu et al., 2005; Anderson et al., 2003; Deschenes-Furry et al., 2007) prompted us to set-up a method (CRAC analysis plus a new dedicated computational pipeline) providing a nucleotide-resolution map of HuD binding in motor neuron-like cells.

HuD has been documented to control the post-transcriptional expression of its target mRNAs, with a described major involvement in mRNA stabilization but with proven roles also in mRNA alternative splicing, alternative polyadenylation, and transport (Bronicki and Jasmin, 2013). For a small set of 6 mRNAs - Nova1, Gls1, Bdnf, Insig1, Atg5, Kv1.1 - HuD binding in the 3’UTR has been associated to enhanced translation (Ince-Dunn et al., 2012; Kim et al., 2014, 2016a; Ratti et al., 2008; Sosanya et al., 2013; Vanevski and Xu, 2015). Interestingly, for 2 other targets - Ins2 and p27 - HuD binding in the 5’UTR resulted in translational repression (Kullmann et al., 2002; Lee et al., 2012). Our collection of HuD binding sites (Figure 2B) shows that in the coding transcriptome HuD is prevalently a 3’UTR binding protein (92% of binding sites).

Functional analysis of HuD interactome revealed, together with the strong neuro-differentiation signature **(Figure 4A)**, another marked functional enrichment which was unexpected. From the CRAC data, HuD resulted to bind up to 80 mRNAs of genes encoding for core components of the general translational machinery, among which 34 ribosomal proteins and 12 translation regulation factors **(Figure 2A)**. This finding prompted us to verify if in our system HuD exerted an effect on global translation. The only available evidence of an action of HuD on global translation comes from the study of the Fujiwara laboratory (Fukao et al., 2009). They demonstrated the binding of HuD to eIF4A1, which results in translation stimulation of a reporter luciferase mRNA in HeLa cell extracts. Interestingly, in their study the presence of the HuD binding site on the reporter construct does not influence translational stimulation, suggesting that indirect effects could be involved. We show for the first time a strong stimulation of HuD on global translation in motor neuron cells, assessed both by the change in polysome formation and by the change in de novo protein synthesis (Figure 2C-D). This global translation enhancement could be likely at least partially mediated by the direct effect of HuD on the initiation factors Eif4A1 and Eif4A2 (Figure 2G). Increased availability of the helicase proteins and the induced HuD overexpression could favour the formation of more HuD/eIF4A complexes (Fukao et al., 2009, 2014), generating a positive feedback loop and boosting HuD effects.

To our knowledge, such an extent of translational stimulation in mammalian cells is only possible by the engagement of the mTORC1 pathway (Saxton and Sabatini, 2017), which mainly targets TOP and TOP-like mRNAs (Meyuhas and Kahan, 2015). Therefore, we checked the degree of coincidence between mTOR responsive genes and HuD targets. We compared experimentally derived sets of mTORC1 responsive mRNAs with HuD translation-related targets. The enrichments reported in Figure 2E clearly demonstrate the high overlap among these lists (3.21 Fold Enrichment, 8.68E-11 FDR).

The mTORC1 pathway is activated in neurons by a wide variety of stimuli, ranging from nutrients to neurotrophic factors to neurotransmitters, and assures neuronal growth and activity by promoting differentiation and synaptogenesis (Takei and Nawa, 2014). Similarly to the HuD-induced phenotype in neurons **(Figure 4A-B)**, the control of protein synthesis through mTORC1 is also essential for axonogenesis and dendritogenesis (Takei and Nawa, 2014), and stimulate axon regeneration in the central nervous system (Berry et al., 2016).

Therefore, we wondered if the newly found HuD control of global translation could act through stimulation of the mTORC1 pathway itself or instead follow an independent route. The several experiments we performed to resolve this issue (Figure 3A-E) consistently favored the second possibility, showing that suppression of the mTORC1 translational burst can be rescued by HuD overexpression. Crucially, this effect is preserved when looking at the translation efficiency of seven distinct mTOR-responsive mRNAs. Moreover, the mRNA responds to HuD with increased translation irrespective of the sequence at the 5’end. Another indirect evidence favouring this model is the smaller, even if statistically significant, decrease in the translational output when HuD is silenced compared to when it is overexpressed (Figure 2C-D). This difference supports the idea of an independent additional effect of HuD rather than a functional integration in the mTORC1 machinery; this function becomes evident only after increased availability of HuD. We believe that this is the first demonstrated control of mTORC1-responsive mRNAs spatially segregated from the mRNA 5’end. Indeed, while recent ribosome profiling experiments have confirmed that mTORC1 acts on the targets through 4E-BP proteins (Thoreen et al., 2012), its sequence-specificity should be reasonably conferred by RBPs binding specific cis-acting sequences. Among the candidates, La protein (Cardinali et al., 2003; Crosio et al., 2000) is a proposed positive effector, while LARP1 (Lahr et al., 2017), AUF1 (Kakegawa et al., 2007), TIA-1 and TIAR (Damgaard and Lykke-Andersen, 2011) are negative modulators. As expected, all these RBPs exert their function binding the TOP or TOP-like motifs at the 5’ terminal, while in all our mTOR responsive mRNAs the HuD binding sites are invariably in the 3’UTR.

These results can be interpreted in terms of a synthetic interaction in motor neurons between the mTORC1 pathway and HuD. Synthetic interactions are widely diffused compensatory mechanisms which contribute to cell robustness against genetic or environmental perturbations (Mohammadi et al., 2012). In the case of motor neurons, we could hypothesize the existence of two independent and redundant triggers of the translational machinery. They could target two spatially segregated portions of the same mRNAs through a fail-safe mechanism to assure the correct translational output in highly polarized cells.

A second unexpected finding from our collection of HuD RNA interactions is the specific and extensive association with the Y3 RNA. Y RNAs are a conserved family of abundant small non-coding RNAs (ncRNA), 100 nucleotides long on average. Through the stem region, Y RNAs form a RNP with the core proteins Ro60, a target of autoimmune antibodies in patients with systemic lupus erythematosus, and La protein, another autoantigen in the Sjögren syndrome (Lerner and Steitz, 1981). The interaction with other RBPs in the loop region has been also reported (Köhn et al., 2013). Although Y RNAs have been known for more than three decades, their cellular functions in vertebrates remain elusive. Well documented is their activity in the nucleus as essential factors for the initiation of chromosomal DNA replication (reviewed in (Kowalski and Krude, 2015).

Using a pan-nELAV antiserum for CLIP analysis in human brain tissue, recently Scheckel and collaborators (Scheckel et al., 2016) reported the first evidence of nELAV binding with 320 different Y sequences. So many different interactors are likely due to the existence of one thousand Y retropseudogenes in the human genome, mostly generated after the rodent/primate divergence (Perreault et al., 2005). The cumulative Y/nELAV binding increased in Alzheimer disease brains and in UV stressed neuroblastoma cells (Scheckel et al., 2016). Our data in murine motor neuron-like cells and with the specific nELAV HuD are instead in favour of a very specific interaction with the Y3 RNA, and always in two positions of the Y3 loop region, fitting the sequence consensus we found for HuD binding (Figure 5A-B). This high selectivity could have been favoured also by the existence in the mouse genome of only 60 Y retropseudogenes, diverged in sequence from the 2 canonical Y RNA genes (Perreault et al., 2007).

Surprisingly, the extent of association between HuD and Y3 in our culture conditions is higher that the cumulative association of the other 1304 coding and 130 non coding RNAs. Considering our estimation of the number of HuD and Y3 molecules per cell (Figure 5C) and assuming the same binding constant between Y3 and non-Y3 HuD binding sites, in our conditions the majority of the expressed Y3 RNA could be associated to HuD. This estimation is instrumental to the hypothesis that Y3 could efficiently modulate HuD in its function as translational enhancer. The subsequent set of experiments convinced us that Y3 negatively affects HuD translational activity by efficiently sequestering it from the translational compartment. Both HuD (Hayashi et al., 2015; Kasashima et al., 1999) and Y3 (Gendron et al., 2001; Peek et al., 1993) are known to be mostly cytoplasmic molecules, but Y3 does not colocalize with polysomes and Y3 silencing induced an increase of HuD polysomal localization, accompanied by improved association to three of its target mRNAs (Figure 5D-E). On the functional side, Y3 depletion increased HuD ability to boost global and transcript-specific translation and effectively rescued HuD depletion (Figure 6A-D). Despite the described antagonism of Y3 and HuD, the fact that La protein also binds Y3 (Lerner and Steitz, 1981) and favours translation of mTOR-responsive mRNAs (Crosio et al., 2000) does not exclude that La could be sponged by Y3 as well, with the same type of effects on translation.

Neuronal differentiation, a measurable cell phenotypic outcome of HuD, was also incremented by Y3 depletion. Finally, we observed a variation of HuD/Y3 level ratio during neural mouse embryo stem cells differentiation (Figure 7A-B). We therefore suggest that a developmentally regulated switch in the HuD/Y3 ratio in vivo may induce release of active HuD, thus boosting neuronal differentiation in a specific temporal window. Interestingly, we also report a localization enrichment of Y3 in mouse primary motor neuron processes, mainly in the axons (Figure S5). HuD has been described to localize in axons and dendrites (Aranda-Abreu et al., 1999; Aronov et al., 2002), and to actively associate with polysomes upon external stimuli, e.g. KCl depolarization (Tiruchinapalli et al., 2008). Therefore, the formation of a HuD/Y3 RNP could in principle contribute to HuD silencing during neuritic transport, triggering translation in neuron microdomains following specific stimuli. Further experiments will be necessary to test this idea; for example it would be interesting to assess if methylation by the CARM1 arginine methylase, known to suppress HuD function (Fujiwara et al., 2006; Hubers et al., 2011), affects HuD binding by Y3.

Our description of an efficient decoy activity on HuD function by Y3 suggests a new role for the Y non coding RNAs, which could extend to other RBPs binding the loop region. The concept of competing endogenous RNAs (Tay et al., 2014) is well established, and applies mostly to microRNAs sequestered from target mRNAs. Functional sequestration of RBPs has been instead described for some lncRNAs (Morriss and Cooper, 2017). One of these lncRNA is cyrano, that sponges the HuD paralogue HuR (Kim et al., 2016b), showed previously by us to associate to Y3 (Köhn et al., 2015). Function of the ELAV RBPs could therefore be controlled by an extensive network of small and long ncRNAs in different cell types.

In conclusion, our work introduces a novel key function for HuD which, besides better describing its cellular effects, could be exploited for therapeutic purposes. Indeed, several evidences suggest the curative potential of positive modulation of the mTORC1 pathway. Limiting to motor neuron diseases, in SMA mice increased translation of the mTOR kinase (Kye et al., 2014) or increased mTORC1 signalling by downregulation of its negative controller PTEN (Ning et al., 2010) rescue axonal defects and improve mice survival. Moreover, in SOD1 mutated ALS mice inhibition of mTOR induces an acceleration of disease signs in vulnerable motor neurons (Saxena et al., 2013; Zhang et al., 2011). For these reasons, attempts aimed at stimulating the mTORC1 pathway, for example with PTEN inhibitors (Chen et al., 2016; Walker and Xu, 2014) could have therapeutic potential for degenerating motor neurons. The new HuD activity we described here of mTORC1-independent global translational enhancer offers another window of opportunity, which becomes even more interesting under the light of the high modulation of HuD function offered by the sponge effect of the ncRNA Y3.

## Acknowledgements

We thank Toshinobu Fujiwara for providing mutated HuD plasmids and Claudia Fallini for supplying the EGFP-HuD vector. We also acknowledge the following CIBIO core facilities for the support: High Throughput Screening, Next Generation Sequencing, Model Organism and Advanced Imaging. We finally thank Viktoryia Sidarovich for valuable suggestions, Paolo Struffi for producing the HuD recombinant protein and Alessandro Roncador for supporting our work with primary neuronal culture. This work was supported by Provincia Autonoma di Trento, (AxonomiX research Project) and by Fondazione Cassa di Risparmio di Trento e Rovereto, Italy. In addition, we acknowledge financial support from the Wellcome Trust funding [109916] and the DFG-funded grant Hu1547/9-1, part of the SPP1935

## Author Contributions

P.Z., T.T. D.P. and A.Q. conceived the study, P.Z., T.T., D.P., M.K.,L.G. and A.Q. designed the experiments, P.Z., D.P., M.K., L.G., V.P. and T.D. performed the experiments, P.Z., T.T., D.P. and G.S. analyzed the data, M.K., S.H., L.C., D.T. P.M. discussed the results and edited the paper, P.Z., T.T. D.P. and A.Q. wrote the paper.

## Supplemental Information

**Supplemental File 1**: collection of 5153 high confidence HuD binding sites identified by CRAC

## Experimental procedures

### Plasmids

To generate pCMV6-HIS-HA-HuD plasmid, the cDNA sequence of human HuD was amplified from SK-N-BE(2) neuroblastoma cell line using the following primers containing Sgf I and Mlu I restriction sites:

Fw HuD 5’-GAGGCGATCGCCGAGCCTCAGGTGTCAAATGG-3′
Rv HuD 5’-GCGACGCGTTCAGGACTTGTGGGCTTTGTTGG-3′

The amplified fragment was digested with Sgfi and MluI enzymes and cloned into the same sites of pCMV6-AN-His-HA vector, that contains an amino-terminal polyhistidine (His) tag and an hemagglutinin (HA) epitope (PS100017, OriGene, Rockville, MD). To generate a lentiviral vector expressing tagged HuD, His-Ha-HuD was excised from pCMV6-AN-His-HA using BamHI and XhoI enzymes and subcloned in the same sites of pENTR-DsRed2 N1 (CMB1) vector. This plasmid was then recombined into pLenti CMV/TO Puro DEST (670-1, Addgene) destination vector using the Gateway^®^ system (Life technologies).

For HuD knockdown, the following oligonucleotides were synthesized and annealed:

5’-GATCCCGCATCCTGGTTGATCAAGTGTGTGCTGTCCACTTGATCAACCAGGATGCTTTTTGGAAA-3’;
5’-AGCTTTTCCAAAAAGCATCCTGGTTGATCAAGTGGACAGCACACACTTGATCAACCAGGATGCGG-3’.

Annealed fragments were ligated into the BglII and HindIII sites of pENTR/pSUPER+ (Addgene 575-1) and transferred into pCMV-GFP-DEST (Addgene 736-1), taking advantage of Gateway technology.

For knockdowns of Y3 by shRNAs, the following oligonucleotides were used:

5’-GATCCCCAACtAAttGAtCACAACCAGtTTCAAGAGAACTGGTTGTGATCAATTAGTTTTTTC-3’
5’-TCGAGAAAAAACTAATTGATCACAACCAGTTCTCTTGAAaCTGGTTGTGaTCaaTTaGTTGGG-3’.

Annealed primers were ligated into pSuperior-GFP (OligoEngine), which was cut with BglII/XhoI. The empty vector served as negative control.

For Y3 overexpression a pGEM-T clone including the whole human Y3 gene was used (Köhn et al., 2015).

To characterize Y3 binding with HuD, the following plasmids, kindly provided by Dr. Toshinobu Fujiwara, were used: pHuD-wt expressing murine HuD wild type (wt), HuD-MUT vector lacking any RNA-binding activity, the HuD-14-302 lacking the poly(A)-binding domain RRM3 and theHuD-216-385 lacking the ARE-binding domain (RRM1 and RRM2).

The luciferase reporter vectors were generated by cloning the specific 3’UTR sequences into *pIS1*-Eef25UTR-renilla vector (Addgene 38235), that harbors a canonic TOP motif in 5’UTR. Specifically, the 3’UTR of Eef1a1, Eif4a1, Eif4a2, Eif4a3 and Rpl10 were amplified from murine cDNA by using the following primers:

Eef1a1 Fw 5’-GCACGGATATCATATTACCCCTAACACCTGC-3’ Rv 5’-GCACGTCTAGACAGATTTCTCATTAAACTTG-3’;
Eif4a1 Fw 5’-GCACGGATATCGGGGCTGTCCTGCGACCTGGCC-3’ Rv 5’-GCACGTCTAGAAGGCAGTTTCCAAGTAATTTTA-3’;
Eif4a2 Fw 5’-GCACGGATATCGGATGAGATAGTTTTGAATGC-3’ Rv5’-GCACGTCTAGACTTCATTAAGACATGTGCAAT-3’;
Eif4a3 Fw 5’-GCACGGATATCAGCTGGTGCTGGTGCACCGAG-3’ Rv 5’-GCACGTCTAGATCACAGGAAAATGTCCACGTT-3’;
Rpl10a Fw 5’-TTTTTGATATCCACGTGAAGATGACCGATGAT-3’ Rv 5’-TTTTTTCTAGAGAGTGGCAGCAGTGAGGTTTAT-3’.

The amplified 3’UTRs were then digested with EcoRV and XbaI enzymes and cloned in the same sites of *pIS1*-Eef25UTR-renilla vector. In addition, Eef1a1 3’UTR and Eif4a3 3’UTR were cloned into pIS1-Eef25UTR-TOPmut-renilla vector (Addgene 38236), that contains a mutated TOP motif in 5’UTR. All plasmids were sequence-verified.

### Cell lines and primary culture

Tetracycline (Tet) inducible cell lines were generated as previously described (Sanna et al., 2015). Briefly, NSC-34 cells were primarily transduced with the pLentiCMV_TetR_Blast vector (716-1, Addgene). To establish an inducible cell line overexpressing the human HuD protein, NSC-34-Trex cells were infected with a lentiviral vector expressing His-HA tagged HuD. Alternatively, NSC-34-Trex cells were stably transfected with pSUPERIOR.neo+GFP plasmid containing the short hairpin sequence for Y3 or the empty vector as a negative control. In both cell lines, the inducible expression of the transgene (HuD or shRNA respectively) was induced by adding 2 μg/ml doxycycline (Clontech) to the culture medium.

Murine NSC-34 and human embryonic kidney HEK-293 cells were cultured in DMEM medium with 10% FBS, 100 U/ml penicillin streptomycin and 0.01 mM l-glutamine (all medium ingredients were obtained from Gibco). Mouse 46C ESCs were cultured in feeder-free conditions as previously described (Ying et al., 2003). Cultures were maintained at 37 °C in a 5% CO_2_ incubator. Primary motor neurons were isolated from embryonic mouse spinal cord and cultured as previously reported (Conrad et al., 2011).To separate motor neuron axons from cell soma and dendrites, the use of coated filter insert (3.0 μm pores PET membrane) was adopted (Poon et al., 2006). After 5 days, the different cellular compartment were rapidly collected by scraping the both sides of PET membranes and RNA was extracted by Trizol (Life Technologies). To qualitatively analyze the separation of motor neuron axons from cell soma and dendritic tree, a small PET membrane piece was cut away, immersed in 4% PFA and processed for immunofluorescence.

### Small interfering RNA (siRNAs) and cell transfections

For gene silencing of Y3, the following siRNA duplexes were used: AACUAAUUGAUCACAACCAGU for Y3 (Köhn et al., 2015)and AGGUAGUGUAAUCGCCUUG as non specific control (47% GC)(Eurofins Genomics); HuD was silenced by transfection of HuD siRNA (sc-37836, Santa Cruz) or control siRNA (sc-37007) from Santa Cruz (Kang et al., 2014). Cells were transfected with 100 nM of the indicated siRNAs for 24h by using Lipofectamine RNAiMAX Reagent (Life technologies). HEK-293 cells were transiently transfected with His-HA HuD plasmid or the His-HA empty vector as control. The transfections were performed using Lipofectamine 2000 (Life Technologies); 48h after transfections cells were harvested for the analysis.

For luciferase assay, HEK-293 cells were transfected with His-HA HuD or His-HA empty vector (75ng). After 24 h, the cells were transfected with both the different renilla luciferase reporter vectors (50 ng) and Firefly luciferase (5 ng) for the normalisation. The luciferase activity was measured after 24h using Dual-Glo^®^ Luciferase Assay System (Promega) following the manufacturer's instructions.

Motor neurons (2 DIV) were transfected by magnetofection using NeuroMag (OZ Bioscience) according to the manufacturer's protocol. At 5 DIV, neurons were fixed in 4% PFA and immunostained.

### NSC-34 cell treatments

To inhibit mTORC1 pathway, NSC-34 cells were starved by serum depletion in DMEM medium without FBS for 8h or treated with Torin1 (500nM) for 1h. After the incubation time, the cells were collected for the following analysis. For the induction of cytoplasmic stress granules, NSC-34 cells were starved for 8h and then treated with 0.25 mM of sodium arsenite (Sigma-Aldrich). After 45 min, the cells were fixed und subjected to immunofluorescence analysis.

### Cell Differentiation

NSC-34 cells were differentiated as reported in (Maier O. et al., 2013). Cells were seeded onto collagen coated (50 ug/mL) 96-well microplate. The normal medium was exchanged 24h after seeding to differentiation medium containing 1:1 DMEM/F-12, 1% FBS, 1% modified Eagle’s medium nonessential amino acids (NEAA), 1% P/S, 5 μM retinoic acid for seven days. HuD overexpression or Y3 short hairpin were induced by 2 ug/mL doxycycline. The differentiation medium was changed after three days.

ESCs were differentiated into neural progenitors and neurons as previously described (Ying et al., 2003). Briefly, dissociated mESCs were plated onto 0.1% gelatin-coated tissue culture dishes at a density of 10^4^ cells per cm^2^ in N2B27 medium. Medium was changed every 1-2 days. N2B27 medium was a 1:1 mix of N2-supplemented DMEM/F12 and B27-supplemented Neurobasal medium (Thermo Fisher).

### Immunofluorescence microscopy

Immunofluorescence of both motor neurons, NSC-34 cells and differentiating ESCs, was performed with the same protocol. After fixation in 4% PFA, cells were permeabilized in PBS + 0.1% Triton X-100 for 5 min and incubated in blocking solution (2% bovine serum albumin, 2% fetal bovine serum, 0.2% gelatin in PBS) for 30 min at RT. Primary antibodies were incubated for 2 hours at RT in blocking solution diluted 1:10 in PBS. The following primary antibodies were used: mouse anti-MAP2 1:300 (M4403, Sigma-Aldrich), rabbit anti-Tau 1:300 (314 002, SynapticSystem), anti-SMI32 (200 KDa neurofilament) 1:300 (Ab7795, Abcam), rabbit anti-MNX1 1:100 (ABN174, Millipore), mouse anti-HuD 1:200 (sc-28299, Santa Cruz), rabbit anti-Tubulin (sc-53140, Santa Cruz), rabbit anti-eEF1A1 (05235, Millipore), rabbit anti-eIF4A2 (31218, Abcam), rabbit anti-PABP 1:500 (Ab21060, Abcam), mouse anti-DCP1A 1:200 (Ab57654, Abcam), goat anti-TIA1 1:100 (sc-166247, Santa Cruz), mouse anti-Oct4 1:400 (sc-5279, Santa Cruz), mouse anti Nestin 1:400 (MAB353, Merck-Millipore), mouse anti beta3-Tubulin 1:1000 (G712A, Promega). The following secondary antibodies, diluted 1:800, were used: goat anti-rabbit Alexa Fluor^®^ 488 (A11008, Thermo Fisher Scientific), goat anti-rabbit Alexa Fluor^®^ 594 (A11012, Thermo Fisher Scientific), goat anti-mouse Alexa Fluor^®^ 488 (A11017, Thermo Fisher Scientific), goat anti-mouse Alexa Fluor^®^ 594 (A11020, Thermo Fisher Scientific), donkey anti-rabbit Alexa Fluor^®^ 488 (A21206, Thermo Fisher Scientific), donkey anti-goat Alexa Fluor^®^ 594 (Ab150136, Abcam). Nuclei were stained with DAPI. Images were acquired with Zeiss Observer Z.1 Microscope implemented with the Zeiss ApoTome device. The objective used for image acquisition was either PlanApo oil immersion lens 63×/1.4 or EC Plan-Neofluor 20x/0.5. Pictures were acquired using AxioVision imaging software package (Zeiss) and assembled with Adobe Photoshop 7.0. Images were not modified other than adjustments of levels, brightness and magnification.

### Neurite outgrowth analysis

NSC34 cells were fixed after seven days of differentiation and stained with Hoechst and mouse anti-Tubulin antibody (1:800; sc-53140, Santa Cruz). For HuD overexpressing cells, an additional immunostaining with an rabbit anti-HA antibody (1:600; A190-1081, Bethyl Laboratories) was performed. The following secondary antibodies, diluted 1:800, were then used: goat anti-rabbit Alexa Fluor^®^ 594 (A11012, Thermo Fisher Scientific), goat anti-mouse Alexa Fluor^®^ 488 (A11017, Thermo Fisher Scientific) and goat anti-mouse Alexa Fluor^®^ 594 (A11020, Thermo Fisher Scientific).Neurite outgrowth was then analyzed on tubulin positive cells by High Content Screening System Operetta™ (PerkinElmer). Briefly, plates (96-well CellCarrier, PerkinElmer) were imaged and acquired in preselected fields with LWD 20x objective. For the feature extraction, the images were analyzed by Harmony software version 4.1 (PerkinElmer). Based on the Hoechst dye cell nuclei were identified. Starting from the cell body region, neurites were then detected in tubulin positive cells. The building block “Find Neurites” automatically calculated for each cell a set of neurite properties.

### CRAC

The CRAC protocol was modified from the published one used in Helwak et al., 2013. Trex-HuD NSC-34 and control Trex NSC-34 cells were seeded onto 150 mm plates (Nunc, Thermo Scientific). Cells were then induced for human HuD production with 10 μg/ml Tetracycline. 24 h post induction growing cells were UV crosslinked on ice with λ = 254 nm in Stratalinker 1800 (Stratagene). The cells were lysed and treated with DNase (Promega M610ACell lysates were incubated with HA agarose beads (26181 Pierce). Ribonucleoprotein complexes on HA beads were trimmed with 0.5 unit RNaseA+T1 mix (RNace-IT, Stratagene 400720-81) and HuD-RNA complexes were eluted. The eluate was incubated with Ni-NTA Agarose (Ni-NTA Superflow 50% suspension IBA 2-3206-010). RNAs bound to HuD were radiolabelled with 32P-γ-ATP and3′ miRCat-33 linker ligation was performed. Then RNA ligase 1 and barcoded 5′ linker were added and the reaction mixture HuD-RNA complexes were eluted by incubation with NuPage-Eluition buffer. Protein-RNA complexes were resolved on a 4%-12% Bis-TrisNuPAGE gel (Life Technologies, NP0335) in NuPAGE SDS MOPS running buffer (Life Technologies, NP0001) and transferred to nitrocellulose membrane (GE Healthcare, AmershamHybond ECL). Air-dried membrane was exposed on film o.n. and the radioactive bands corresponding to the HuD complexes were cut out. RNA was extracted and reverse transcribed. cDNA was amplified and PCR products were precipitated, resuspended and separated on a 2.5% MetaPhoragarose (Lonza). After purification with Gel Extraction Kit with MinElute columns (QIAGEN) the samples were sequenced on Illumina platform.

### Polysome profiling

Polysomal profiling was performed according to previously described protocols (Dassi et al., 2013; Viero et al., 2015). Briefly, the cells were treated with cycloheximide and then lysed in 300μl of cold lysis buffer. The lysate was centrifuged at 4 °C for 5min at 20.000 g to pellet tissue debris. The cytoplasmic lysates loaded on a linear 15–50% [w/v] sucrose gradient and centrifuged in a SW41Ti rotor (Beckman) for 1 h 40 min at 180.000 g at 4 °C in a Beckman Optima^TM^ Optima XPN-100 Ultracentrifuge. Fractions of 1 ml of volume, were then collected monitoring the absorbance at 254 nm with the UA-6 UV/VIS detector (Teledyne Isco).

### Extraction of total and polysomal RNA

Sucrose fractions corresponding to polysomes and total RNA were pooled together and the RNA was extracted with phenol/chloroform. Purity of RNAs and concentration was measured using Nanodrop spectrophotometer. The total RNAs were isolated using Trizol reagent (Life Technologies) following the manufacturer’s instructions.

### RT-qPCR analysis

The retrotranscription reaction was performed with 1 μg of polysomal or total RNA using the iScriptcDNA synthesis kit (Biorad) in accordance with the manufacturer's instructions. The obtained cDNA was used as template in aqPCR reaction with the KAPA SYBR FAST qPCR (Kapa Biosystem) and specific primers as reported in the following **Table of the primers**:

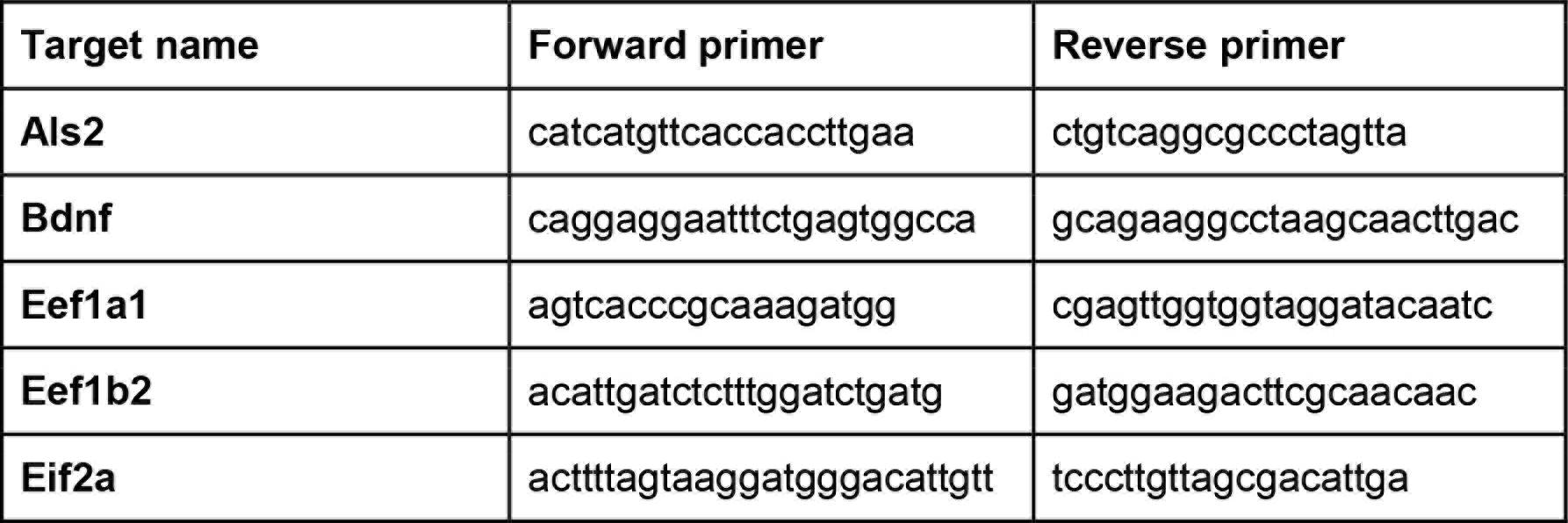

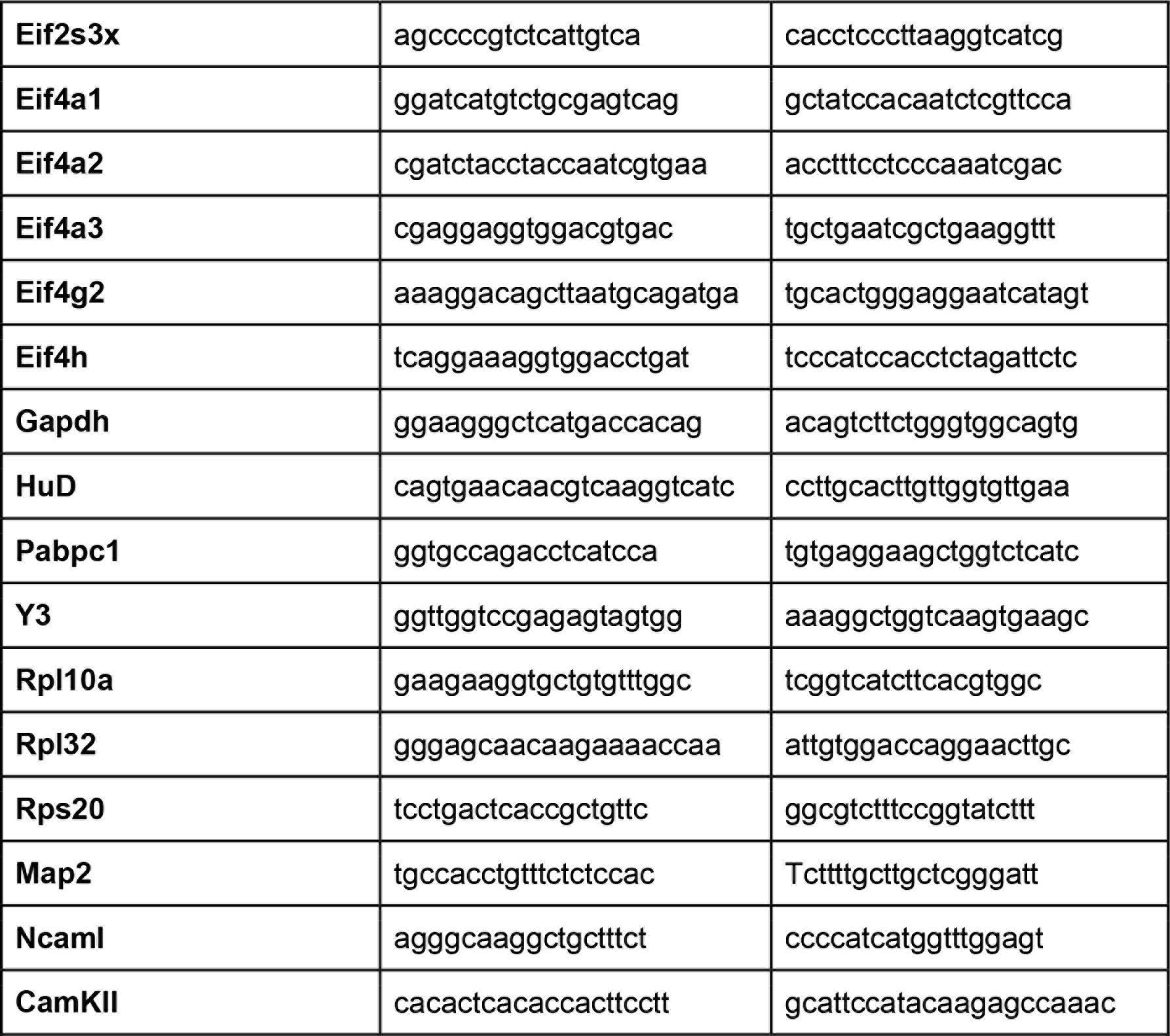

qPCR were run in three biological and three technical replicates. The relative expression was calculated with the delta delta Ct method. Gapdh and Als2 were used as reference genes. The gene-specific Translation Efficiency (TE) was calculated as the ratio between the fold change at the polysomal level and the fold change at the total level of the gene of interest.

### TruSeq^®^ Targeted RNA Expression Library preparation

The library was prepared using TruSeq^®^ Targeted RNA Expression following the manufacturer’s instruction and the sequencing was performed on the MiSeq Illumina platform.

### RNP immunoprecipitationand RNA pulldown

The HuD ribonucleoproteincomplex was isolated as previously described (Sanna et al., 2015). Immunoprecipitated and input samples were resuspended in Trizol reagent (Life Technologies) and RNA extraction was performed following manufacturer’s instructions.

RNA pulldowns were essentially performed as previously described (Köhn et al., 2015). For synthesis of the RNA baits (Y1, Y3, Y4) T7-Polymerase mediated *in vitro* transcription was used.

### SBP Pulldown

To pull down HuD and HuD-fragments inserts were cloned into the pCDNA-SBP-Flag vector. After transfection into NSC-34, cells were harvested after 48h. Cell pellets were lysed using BB (100 mMKCl, 10mM EDTA, 10 mM HEPES pH 7.4, 0.5% NP-40) and the supernatant was incubated with Streptavidin MyOne T1 beads (Life Technologies). Beads were then washed three times with BB and bound proteins were eluted by addition of BB+1% SDS and heating at 65°C. Eluates were then separated for RNA and protein preparations. Input and pulldown RNA was purified using Trizol (Sigma-Aldrich) and subjected to Northern Blot. Protein samples were subjected to Western Blot.

### Northern and Western blot

Northern Blot was essentially performed as previously described (Köhn et al., 2013, 2015).For Western blot analysis, NSC-34 and HEK-293 cells were homogenized in RIPA lysis buffer (SIGMA) following the manufacturer’s instructions. Protein lysates (30 μg) were resolved on SDS-PAGE and transferred to nitrocellulose membrane. The following antibodies were used: mouse anti HuD (sc-28299, Santa Cruz), rabbit anti HA (A190-1081, Bethyl Laboratories), rabbit anti eIF4A1 (ab312-17, Abcam), rabbit anti-eIF4A2 (31218, Abcam), rabbit anti-eIF4A3 (homemade, generously provided from Prof. Macchi's lab), rabbit anti eEF1A1 (SAB2108050, Sigma), Tubulin (sc-53140, Santa Cruz).

### Quantification of HuD and Y3 molecules

NSC-34 cells (5×10^6^) were lysed using RIPA buffer and the protein concentration was determined using standard Bradford Protein assay (Sigma). Known amount of cell lysates and HuD recombinant protein (a generous gift of Dr. Paolo Struffi, University of Trento) were separated by 10% SDS-PAGE and transferred to nitrocellulose membrane. Samples were analyzed by western blotting using rabbit anti-HuD antibody (sc-28299, Santa Cruz) and the optical density (OD) of the protein bands were quantified by ImageJ. To estimate the number of HuD molecules NSC-34 cells, a standard curve was generated by plotting the known amounts of HuD recombinant protein (15, 25, 50, 75 ng) on the X axis, and their respective OD values on the Y axis. This reference plot was used to inferred the amount of HuD protein in our NSC-34 lysate and calculate the amount for cell.

To estimate the number of Y3 molecules in NSC-34 cells, murine Y3 was synthesized by *in vitro* transcription. Total RNA was extracted from NSC-34 cells with Trizol (Sigma-Aldrich). The amount of RNA was normalized to the cell number and corrected for purification efficiencies. Then quantitative Northern Blots were performed to determine the amount of Y3 in NSC-34 total RNA by using *in vitro* transcribed Y3 as a standard. Finally, the amount of Y3 per NSC-34 cell could be determined.

### AHA assay

*De novo* synthesized proteins were quantified using the Click-iT^®^ AHA Alexa Fluor^®^ Protein Synthesis HCS Assay (Molecular Probes, Life Technologies). In brief, NSC-34 cells were plated at density of 10,000 cells/well in 96-well plates for 24h.The cells were then induced to overexpress HuD (a) or silenced for HuD (b) or silenced for Y3 (c) or subjected to HuD overexpression and Y3 silencing (d). After 48h, the cells were washed, incubated with L-azidohomoalanine (AHA) 50 μM for 1h and fixed. During AHA incorporation, control cells were treated with puromycin (100 ug/ml), a protein synthesis inhibitor, to evaluate background labeling. Click-chemistry reactions were sequentially performed according to the manufacturer’s instructions and the relative AHA incorporation was then analyzed by high content imaging approach. To detect cell nuclei, the kit was multiplexed with Hoechst 33342. Plates (96-well CellCarrier, PerkinElmer) were imaged on the High Content Screening System Operetta^TM^ (PerkinElmer). In each well, images were acquired in preselected fields with LWD 20x objective. For the feature extraction, the images were analyzed by Harmony software version 4.1 (PerkinElmer). Based on the Hoechst dye and Alexa 488 fluorescence intensity, cell nuclei and cell cytoplasm were identified respectively. To quantify nascent protein synthesis, the mean fluorescence intensity of Alexa Fluor 488 was quantified in the cytoplasm.

### CRAC data analysis

Adapter removal and collapse of duplicate reads (also with identical random barcode, marking PCR duplicates) were performed with FASTX-Toolkit (0.0.14). Reads were aligned to the mouse genome (GRCm38.p4) with Tophat (version 2.0.14), using the Gencode M6 transcript annotation as transcriptome guide. All programs were used with default settings unless otherwise specified. In order to detect CRAC binding sites, we developed and implemented a dedicated computational methodology (MAPAS, standing for Mutation And PWM Assisted Search) that takes advantage of cross-linking induced mutations, consisting primarily in deletions in our experiment, in order to localize candidate binding sites (similar in this step to CIMS, (Zhang and Darnell, 2011)). After the integration of replicates, to increase specificity, we penalized locations with aligned reads and deletions in control experiments (noise subtraction and removal). For each of the remaining locations, we calculated a combined p-value based on a) the number of deletions, b) the number of aligned reads (coverage). P-values were empirically calculated from the genome-wide experimental distributions of coverage and number of deletions. Coverage and deletion p-values were combined with the Fisher method.

A pool of 753 sequences surrounding unique genomic locations with a combined p-value < 0.05 were selected to build a PWM (Positional Weight Matrix), hence used as a “seed” matrix to score to all the other candidate binding sites. To create the seed PWM, we defined a region spanning seven nucleotides around the deletion site. This size choice is based on previous crystallographic studies resolving the structure of the RRM1 and RRM2 domains of HuD bound to canonical AU rich elements. PWM analysis was performed with functions implemented in the Biostrings R package. The seed PWM was used to score all deletion sites and select high-confidence HuD bound sites. A PWM score threshold was chosen, based on the 95 percentile of scores obtained from random heptamers. HuD “high confidence” binding sites were selected among those with i) PWM score > PWM score threshold, ii) number of HuD deletions >= 3 (at least one for replicate), iii) number of aligned reads >= 6 (at least two for replicate). This procedure identified 5153 high confidence HuD binding sites (Supplemental file 1).

### RNA-Seq data analysis

For RNA-Seq data of NSC-34 cells, after quality control (FastQC) reads generated from each sample were aligned to the mouse genome (GRCm38.p4) with Tophat (version 2.0.14), using the Gencode M6 transcript annotation as transcriptome guide. Mapped reads were subsequently assembled into transcripts guided by reference annotation (Gencode M6) with Cufflinks (version 2.2.1). Expression levels for genes and transcripts were quantified as normalized FPKM (fragments per kilobase of exon per million mapped fragments) using Cufflinks. The experiment was performed in biological duplicate.

### TruSeq^®^ Targeted RNA Expression data analysis

TruSeq Targeted sequencing of 75 genes (including 70 HuD targets and 5 negative control genes) was performed to validate HuD RNA interactome (RIP-Seq in NSC-34 cells) and to monitor expression variations of HuD targets upon HuD overexpression (total RNA and polysomal RNA in NSC-34 cells), with the Illumina MiSeq platform. Raw counts were determined from the alignment of reads to targeted gene sequences. For HuD overexpression assay, normalization with the TMM method and identification of differentially expressed genes (p-value <0.05) were performed with the edgeR package. For the RIP-seq assay, negative control genes were used as housekeeping and data were normalized for the geometric mean of their expression values. Experiments were performed in biological triplicate.

### Functional annotation enrichment analysis

Functional annotation enrichment analysis with Gene Ontology terms, KEGG and REACTOME pathways were performed using the clusterProfiler Bioconductor package.

Functional annotation enrichment analysis with lists of genes derived from experimental datasets was performed with the enrichR gene set libraries.

## Supplementary Figures

**Figure S1. Related to Figure 1.**
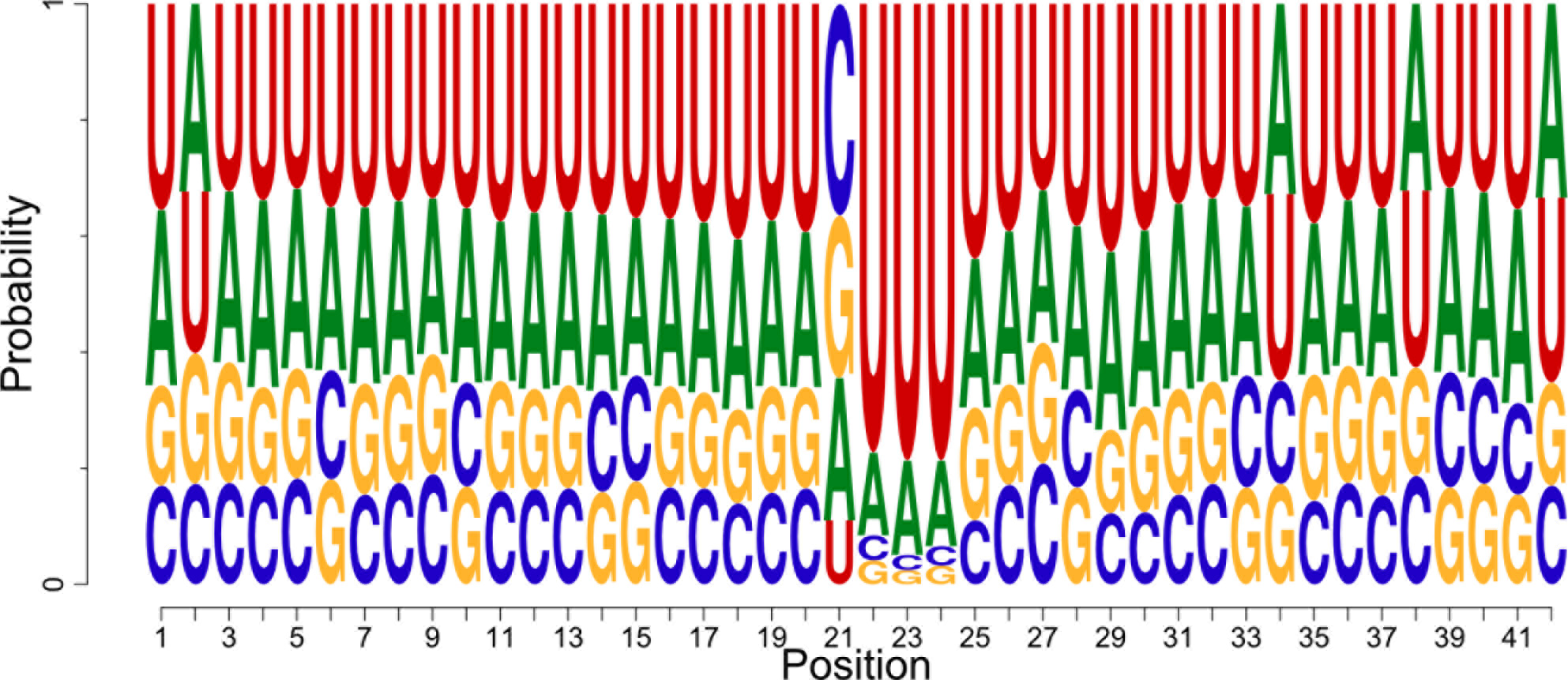
Extended sequence logo determined from CRAC unique deletion locations (position 22).

**Figure S2. Related to Figure 3.**
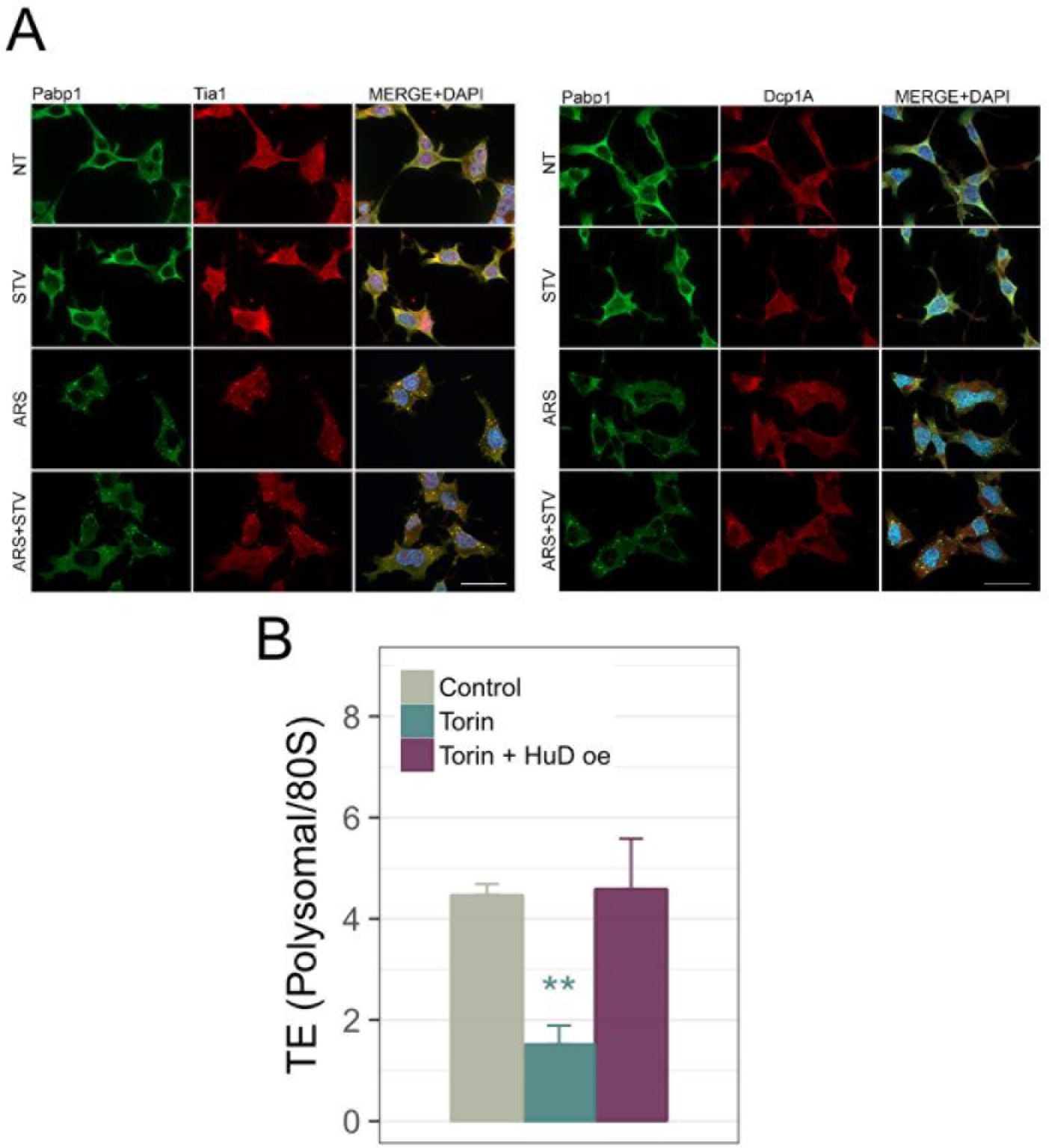
**(A)** Starvation in NSC-34 cells induces neither stress granule nor P-bodies formation. Left panel: NSC-34 were immunostained for Pabp1 (green) and Tia1 (red) to detect stress granules formation. Arsenite treatment was used as positive control for stress granules formation. The images show that starvation per se does not induce stress granules and does not increase arsenite-triggered stress granules. Right panel: NSC-34 were immunostained for Pabp1 (green) and Dcp1a (red) that is a P-bodies marker. Arsenite treatment was used as positive control for increasing P-bodies’ size and number. The images show that starvation does not alter P-bodies in neither size nor number. The scale bars correspond to 40 μm. **(B)** Measurement of global TE by sucrose gradient centrifugation as the ratio between the area under polysome peaks and the area under the 80S peak in the following conditions: control, Torin 1 treatment and Torin 1 treatment coupled with HuD overexpression. (data are represented as mean ± SEM. t-test *p < 0.05, **p < 0.01 and ***p < 0.001 “Torin” was compared to “Control”, and “Torin + HuDoe” was compared to “Torin” for testing statistical significance).

**Figure S3. Related to Figure 4.**
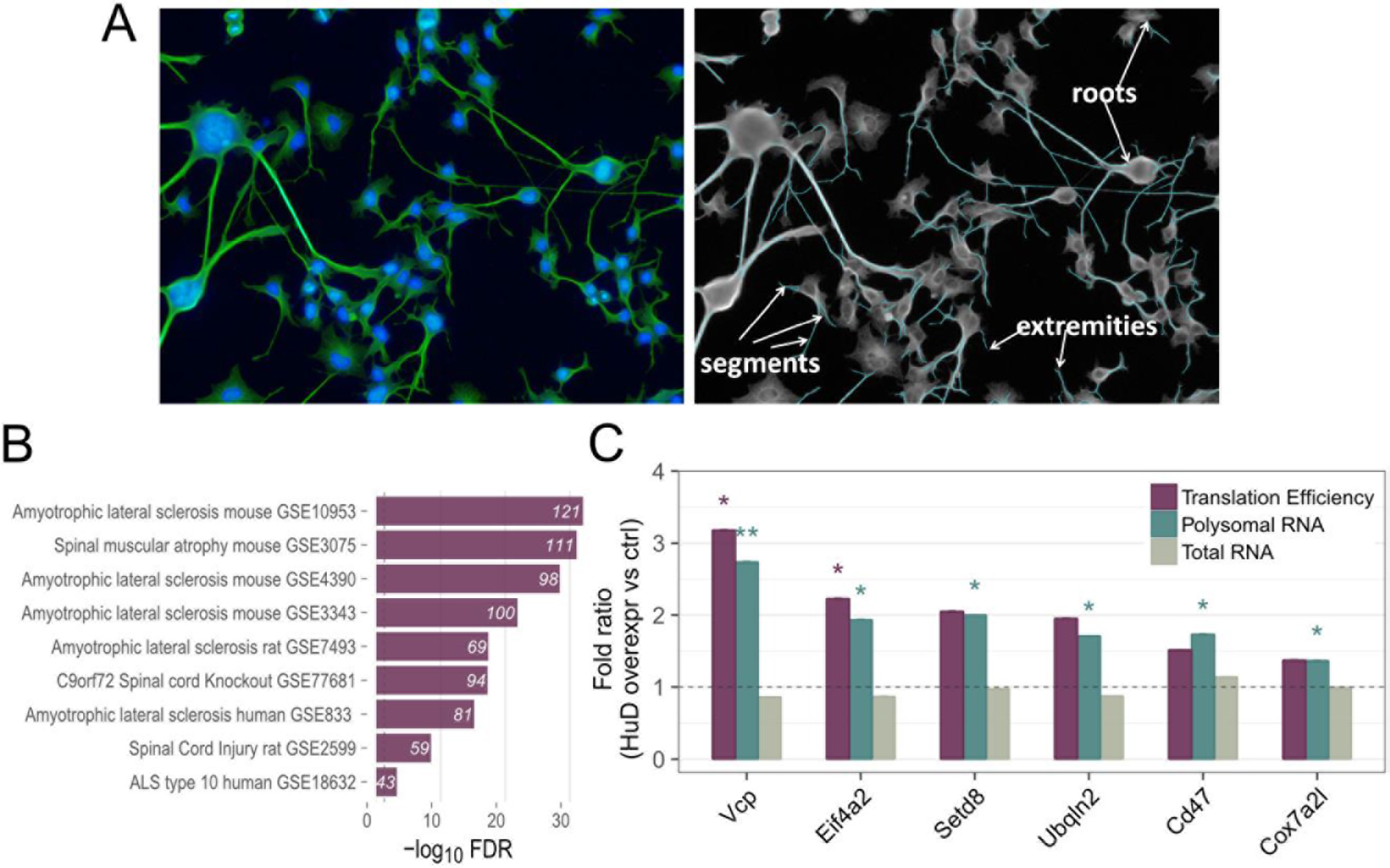
**(A)** Representative images of differentiated NSC-34 cells (left). The cytoplasm was immunostained with anti-tubulin antibody (green) to detect neuritic arborization, whereas the nuclei was identified by Hoechst staining (blue). Automated neurite segmentation performed with an Operetta HCS device (right). The cells were imaged on the High Content Screening System Operetta^TM^ (PerkinElmer) and the number of neurite segments, of extremities and of roots were quantified. (B) Enrichment of HuD targets among collections of genes with altered expression levels in motor neuron diseases (available in enrichR libraries). **(C)** Measurement of TE variations of HuD targets related to motor neuron diseases upon HuD overexpression in NSC-34 cells. TE was calculated as the ratio between the polysomal and total RNA quantified by targeted sequencing and displayed in the plot in green and grey, respectively. *(In panels C data are represented as mean ± SEM. t-test *p < 0.05, **p < 0.01)*.

**Figure S4. Related to Figure 5.**
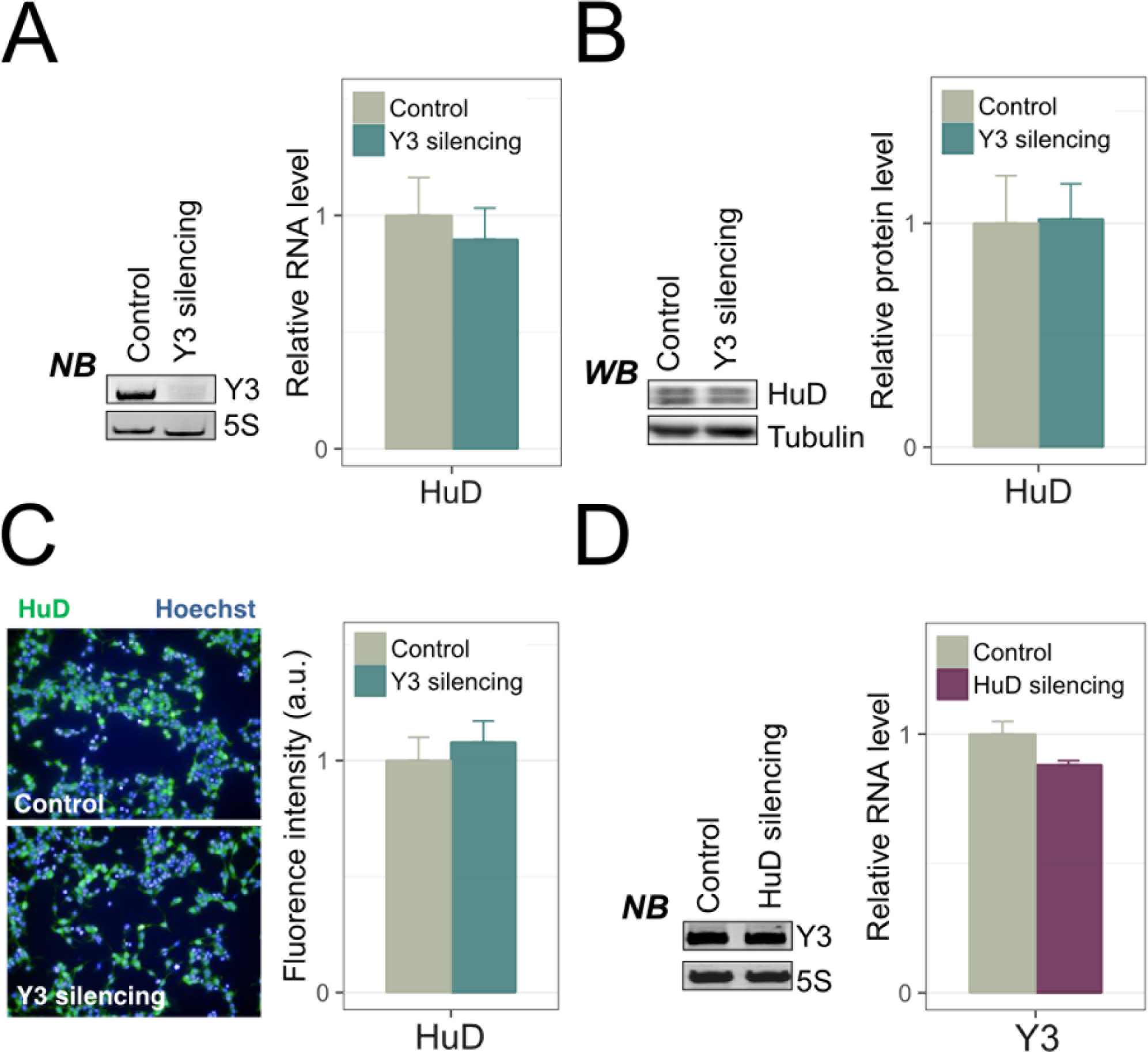
HuD and Y3 do not mutually influence their expression. **(A)** Left: Northern blot showing the efficient silencing of NSC-34 transfected with si-Y3 in comparison with control (scramble). 5S is used as housekeeping; right: level of HuD transcript quantified by RT-qPCR in si-Y3 NSC-34 cells compared to control. Rpl10a and Gapdh were used as reference. **(B)** HuD protein levels quantified by Western Blot in control and Y3-siRNAcells. **(C)** HuD protein levels quantified by immunofluorescence in control and Y3-siRNA cells. **(D)** Y3 quantification by Northern blot in control and si-HuD cells. NB, WB and IF images are representative of at least three biological replicates.

**Figure S5. Related to Figure 6.**
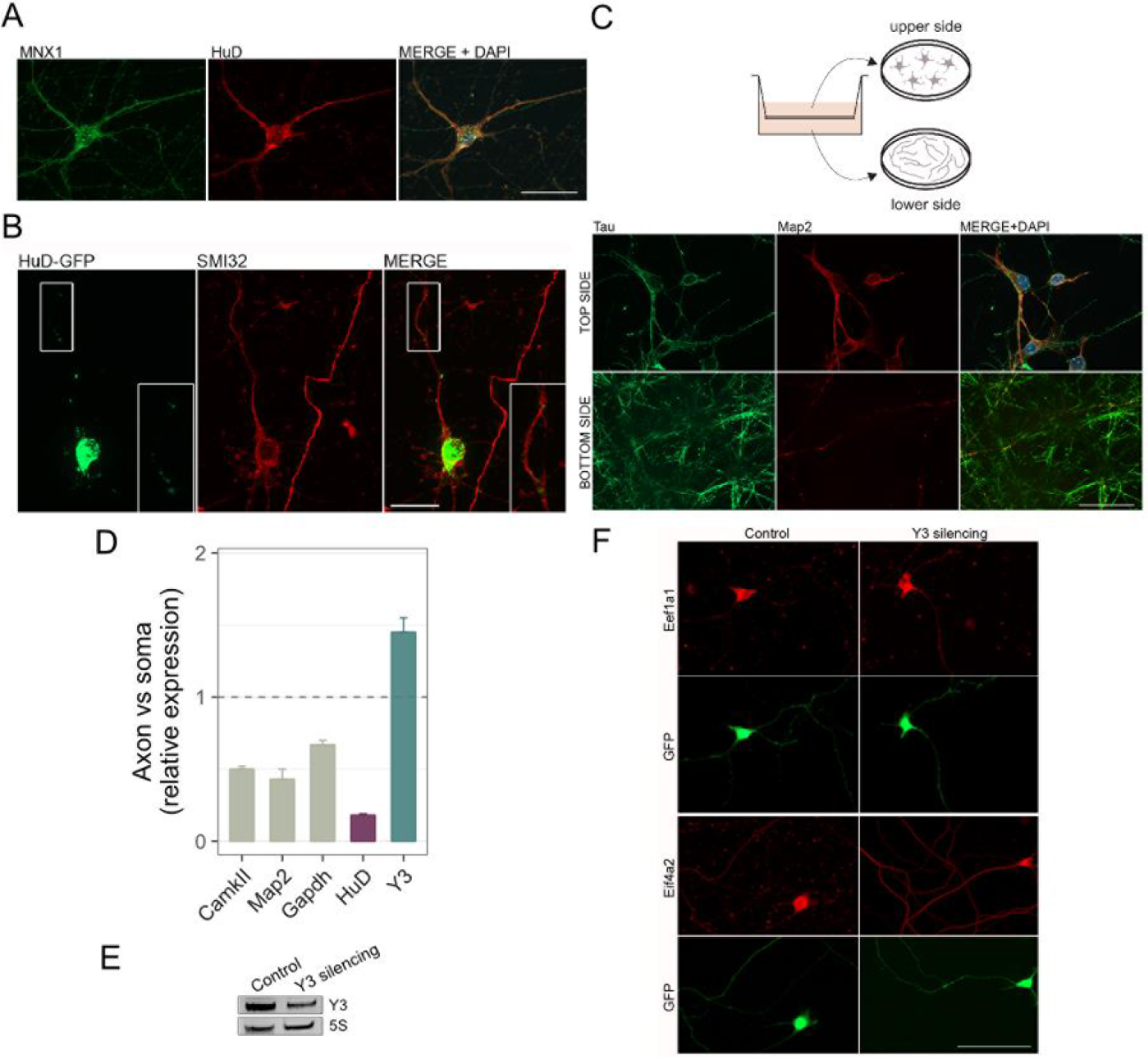
HuD and Y3 are both expressed in motor neurons with a Y3 enrichment in axonal compartment. **(A)** Primary motor neurons (5 DIV) immunostained for the motor neuronal marker Mnx1 (Hb9, green) and HuD (red). The scale bar corresponds to 40 μm. **(B)** Motor neurons immunostained for the motor neuronal marker SMI32 after 48 hours transfection with pHuD-GFP vector. Both endogenous and transfected HuD shows a granular staining in the cell body, dendrites and along the axons. The scale bar corresponds to 20 μm. **(C)** Top: Schematic overview of the separation system. Embryonic motor neurons were cultured in tissue culture inserts fitted with PET membrane with 3 μm pores. Neuron cell bodies remained on the upper membrane surface, whereas axons and few dendrites passed through membrane pores reaching the lower side of the membrane. Bottom: Immunofluorescent staining of motor neurons on PET membrane: neuronal processes were marked with Tau (green), dendrites Map2 (red), cell nuclei with DAPI (blue). Images confirm that mainly axons (just few dendrites) cross the membrane and reaches the bottom side, with no contamination of cell bodies. On the contrary, cell bodies and dendrites were retained on the top side. The scale bar corresponds to 40 μm **(D)** RT-qPCR analysis of RNA extracted from motor neuron axons (scraped from the membrane bottom side) and compared with RNA extracted from cell bodies and dendrites (scraped from the membrane top side). Relative expression data show that *Y3* RNA is more present in the axons compared to cell soma and dendrites, while *HuD* and *Map2* are more localized to cell soma and dendritic tree. **(E)** Assessment of the effective silencing of *Y3* RNA using the Y3-short hairpin vector. **(F)** Immunostaining of Eef1a1 and Eif4a2 protein levels in primary motor neurons transfected with an shRNA construct directed against Y3 (sh_Y3) or a control vector (sh_Ctrl). GFP (green) identifies transfected cells that were subjected to quantification of either Eef1a1 or Eif4a2 signal (red). (n = >20 cells/condition).

## Footnotes

1 Lead contact: Alessandro Quattrone, Laboratory of Translational Genomics, Centre for Integrative Biology, University of Trento, Trento, Italy.

